# Post-Translational Modifications Optimize the Ability of SARS-CoV-2 Spike for Effective Interaction with Host Cell Receptors

**DOI:** 10.1101/2021.12.02.470852

**Authors:** Karan Kapoor, Tianle Chen, Emad Tajkhorshid

## Abstract

SARS-CoV2 spike glycoprotein is prime target for vaccines and for diagnostics and therapeutic antibodies against the virus. While anchored in the viral envelope, for effective virulance, the spike needs to maintain structural flexibility to recognize the host cell surface receptors and bind to them, a property that can heavily hinge upon the dynamics of the unresolved domains, most prominently the stalk. Construction of the complete, membrane-bound spike model and the description of its dynamics remain critical steps in understanding the inner working of this key element in viral infection. Using a hybrid approach, combining homology modeling, protein-protein docking and MD simulations, guided by biochemical and glycomics data, we have developed a full-length, membrane-bound, palmitoylated and fully-glycosylated spike structure in a native membrane. Multi-microsecond MD simulations of this model, the longest known trajectory of the full-spike, reveals conformational dynamics employed by the protein to explore the crowded surface of the host cell. In agreement with cryoEM, three flexiblele hinges in stalk allow for global conformational heterogeneity of spike in the fully-glycosyslated system mediated by glycan-glycan and glycan-lipid interactions. Dynamical range of spike is considerably reduced in its non-glycosylated form, confining the area explored by the spike on the host cell surface. Furthermore, palmitoylation of the membrane domain amplify the local curvature that may prime the fusion. We show that the identified hinge regions are highly conserved in SARS coronaviruses, highlighting their functional importance in enhancing viral infection, and thereby provide novel points for discovery of alternative therapeutics against the virus.

**Significance:** SARS-CoV2 Spike protein, which forms the basis for high pathogenicity and transmissibility of the virus, is also prime target for the development of both diagnostics and vaccines for the debilitating disease caused by the virus. We present a full model of spike methodically crafted and used to study its atomic-level dynamics by multiple-*µ*s simulations. The results shed new light on the impact of posttranslational modifications in the pathogenicity of the virus. We show how glycan-glycan and glycan-lipid interactions broaden the protein’s dynamical range, and thereby, its effective interaction with the surface receptors on the host cell. Palmitoylation of spike membrane domain, on the other hand, results in a unique deformation pattern that might prime the membrane for fusion.

## Introduction

The pandemic caused by SARS-CoV-2 continues to impact nearly all aspects of human lives across the globe.^1, 2^ SARS-CoV-2 belongs to the beta coronavirus genus, same as SARS-CoV and MERS-CoV responsible for the severe acute respiratory syndrome outbreak in 2003 and Middle East respiratory syndrome in 2012, respectively.^3, 4^ The first step in viral entry and infection in SARS-CoV-2, and in other coronaviruses in general, is the binding of its extended spike glycoproteins located on the viral envelope to specific human cell surface receptors.^5, 6^ These envelope spike proteins contain all key components necessary for the infection of human cells; after binding to the surface receptors they initiate the fusion of the viral and human membranes and the subsequent release of the viral genome into the host cell.^7^ Characterizing the mechanism of action of the spike protein thus remains key to our understanding of the critical steps involved in viral infection, paving way for development of novel therapeutics against the virus.

Major efforts have been spent recently in characterizing the structural features of the spike, and as such, quite a few SARS-CoV-2 spike cryo-EM structures have been resolved.^8–15^ Though providing essential information on the structural details of the spike globular domain (spike head), which contains the receptor-binding domain (RBD) recognizing and binding to the ACE2 host cell receptors,^6^ the resolved structures are still missing a number of functionally important regions in the full spike (Fig. 1A). These include the highly conserved fusion-peptide segment involved in initiating the fusion of the viral and host cell membranes,^16, 17^ the stalk heptad repeat 2 (HR2) domain undergoing refolding during the membrane fusion, ^18^ and the transmembrane (TM) domain containing multiple palmitoylation sites shown to stabilize the spike in the viral envelope and thus essential for assembly and activity in other viruses.^19, 20^ The structures also lack critical information about the glycosylation of different residues, known to be essential in mediating protein folding, shaping viral tropism, as well as shielding the virus from immune recognition.^21^ Additionally, the assembly and budding of the virus are known to take place in the endoplasmic reticulum–Golgi intermediate compartment (ERGIC),^22, 23^ an environment that is not taken into account in any of the above cryo-EM studies which have resolved only the ectodomain structure stabilized using C-terminal foldon trimerization motif.^8^ Further resolution of these structural and environmental details thus becomes a necessary step to study the behavior of native spike proteins in a physiological context.

**Figure 1:**
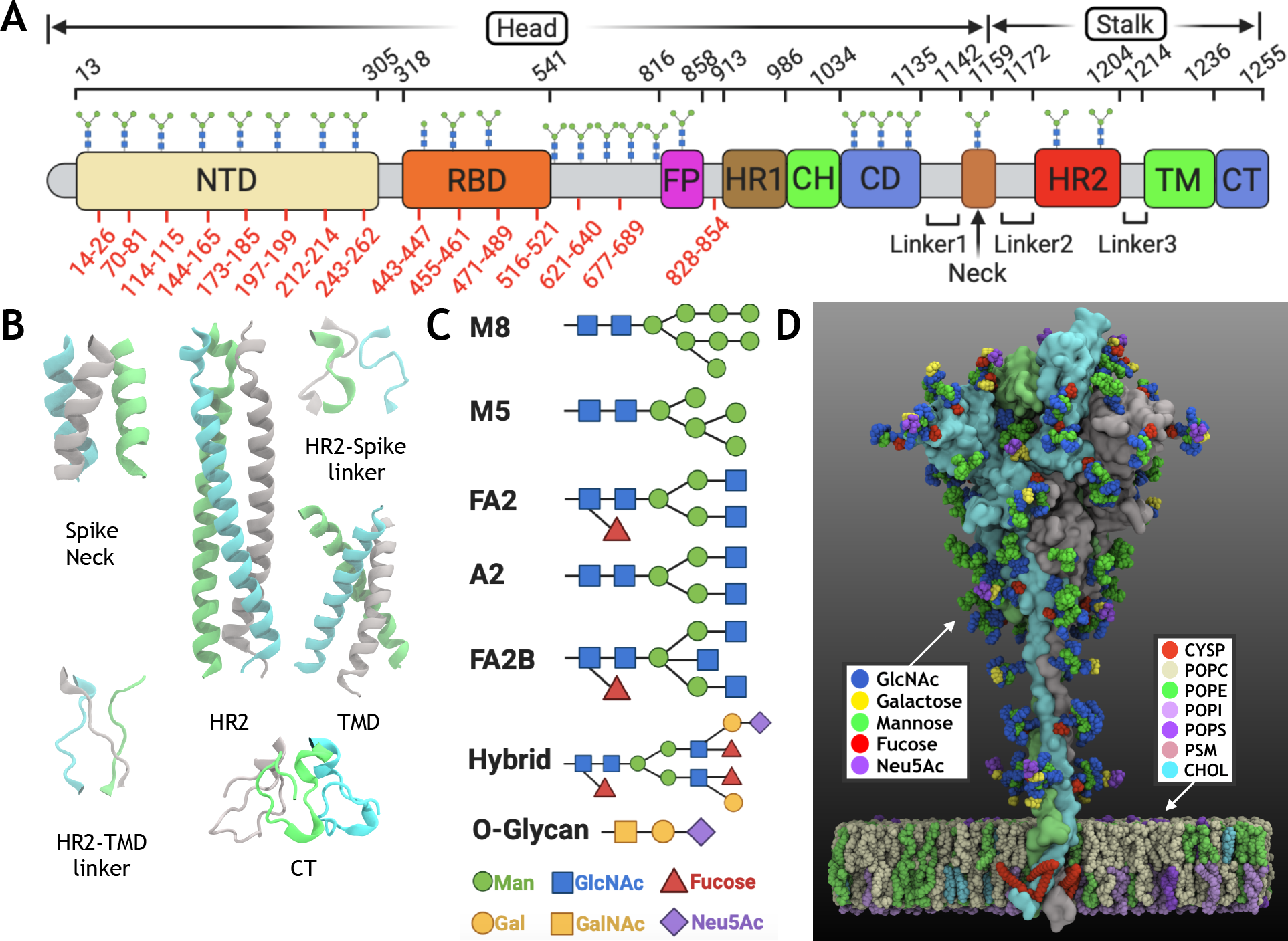
Overview of full spike system: (A) Schematic of the different functional domains of the spike, which can be largely divided into the head and stock domains: N-terminal domain (NTD), receptor binding domain (RBD), fusion peptide (FP), heptad repeat 1 (HR1), central helix (CH), connecting domain (CD), spike neck, heptad repeat 2 (HR2), transmenbrane domain (TM) and cytoplasmic tail (CT). The residue ranges of these domains are provided along with the experimentally identified glycosylation sites marked on top of the domains. Out of these, cryo-EM structures are only available for the spike head, with the missing regions in the structure shown in red. (B) The modeled structures in the spike neck, HR2, TM domain, CT and the 2 connecting linkers are shown. These missing regions in the spike stalk were modeled using a hybrid approach (described in methods). (C) The types of N-linked and O-linked glycans modeled in the full spike structure are shown. (D) The modeled structure of the full-length, palmitoylated and glycosylated spike is shown. Each spike monomer is depicted in surface representation in a different color. The protein is embedded in a membrane bilayer with a lipid composition from endoplasmic reticulum-Golgi intermediate compartment (ERGIC). The constituting glycan components and lipid/palmitoylation types are provided in the legends.

Computational simulations offer an effective technique for modeling complex systems like the spike at a detailed level. As such, recent efforts in applying classical molecular dynamics (MD) simulations to the resolved structures of the spike head^24–27^ as well as constructed full-length spike models^28–30^ have provided powerful insights into various functionally relevant features, e.g., how glycans can shield the spike from the immune system and neutralizing antibodies,^24, 25, 29, 30^ or the intra-domain dynamics of the spike head that may facilitate the RBD to switch between the ’up’ and ’down’ (active or inactive) conformations. ^26, 27, 29^ However, we are still far from fully understanding how the global dynamics of the spike may contribute to its function, specifically in relation to effectively locating its target receptor in crowded cellular environments. So far, characterization of the effect of the spike glycosylation has been focused on their global shielding effects, leaving the possible roles that the glycans may play in the protein’s conformational dynamics unexplored. Furthermore, the role of palmitoylations present in the TM/endodomain, known to participate in the modulation of the membrane curvature and in mediating cell fusion in other viruses,^31–34^ also remains lacking in SARS-CoV-2.

To obtain further insights into the workings of this important component of the virus machinery, we aimed to construct a full-length, membrane-embedded spike for investigation of its conformational dynamics at the atomic-scale using MD simulations. In the first step, we use a hybrid approach, combining homology modeling, protein-protein docking and MD simulations of the modeled regions, with the available experimental data,^35–38^ to develop a full-length, palmitoylated and fully-glycosylated spike. After validating the modeled parts, multi-microsecond MD simulations of the full-length spike, embedded in a lipid bilayer of composition derived from ERGIC, were used to characterize global dynamics and inter-domain motions, focusing specifically on the potential role of glycans in these processes. Control, multi-microsecond simulations of the non-glycosyslated spike highlighted the importance of the glycans in modulating the global motion of the spike. We also characterized the specific role played by palmitoylations in modulating the membrane curvature, potentially regulating the membrane fusion with the host cell. This study provides key insight into the dynamics used by the spike to sample the crowded surface of its host cell and the direct role of the glycans in regulating this process while successfully allowing the virus to evade the host immune response.

## Methods

### Modeling of full-length spike

Topologically, the spike can be divided into two main regions - the spike head, which includes the RBD that binds to the host cell receptor, and an extended stalk consisting of a short spike neck, the HR2 domain, and the TM domain tethering the spike to the viral envelope. In addition, the spike also contains a cysteine-rich, disordered C-terminal region inside the envelope. Since the experimental structures have only been resolved for the spike head, as the first step, we model the missing regions to construct a full-length spike structure, which is then palmitoylated at the TM/endodomain and fully-glycosylated, before performing MD simulations. The different steps involved in this process are described below.

### Constructing missing regions of the spike head

The recent cryo-EM structures of the spike head don’t provide coordinates for a number of missing regions including a short peptide segment called the fusion peptide, known to to be important for membrane binding and insertion,^16, 17^ as well as a number of loops. These missing regions were generated using either template-or fragment-based modeling as described below. Firstly, the cryo-EM structures of the spike head from SARS-CoV-2 (PDB: 6VYB and 6VXX^9^), SARS-CoV (PDB: 5X58,^39^ 5XLR,^40^ 6CRW^41^) and MERS-CoV (PDB: 6Q04^42^), along with the SARS-CoV-2 RBD structure in complex with human receptor ACE2 (PDB: 6M17^43^) were aligned and using sequence-and structure-based alignment in MOE.^44^ The missing regions were then constructed in MOE by grafting a suitable template region from the aligned structures and using local superimposition based on the flanking residues of the template region. Residues in the grafted templates from other CoV structures were mutated back to those in the SARS-CoV-2 sequence. Missing loop regions in the spike head without a suitable template were constructed using Rosetta ab-initio fragment assembly in the Robetta protein structure prediction server.^45, 46^

### Modeling the neck and the HR2 domain

Due to lack of a suitable template, the spike neck was modeled using ab-initio modeling in Robetta. The HR2 domain was homology-modeled using the NMR-resolved structure of homologous SARS-CoV (PDB: 2FXP^47^) as the template in MOE. These modeled regions were validated against secondary structure predictions by JPred4.^48^

### Developing a trimeric TM model

The structural information about the TM region of SARS-CoV-2 spike, which is highly conserved among coronaviruses and known to play a role in the spike assembly and stabilization,^19, 49^ is limited, as no structure for this domain has been resolved. The available structure from the HIV virus displays a low sequence homology, making it unsuitable for template-based modeling.^50^ The TM domian of SARS-CoV-2 was thus built in an ab initio manner. Firstly, the location of the TM region in the spike sequence was predicted using TMHMM,^51^ a bioinformatics-based approach for predicting membrane spanning regions in protein sequences. Using the predicted TM sequence, a TM helix monomer was constructed in MOE. The monomer was then placed in a membrane patch using CHARMM-GUI^52, 53^ with a lipid composition based on the lipoprofiles of the ERGIC,^22, 54^ where coronaviruses assemble.^23^ The specific lipid compositions of the two leaflets are provided in Table S1. After solvating the system, 1,000 minimization steps and 100 ns equilibration were performed under NPT conditions to relax the monomeric structure in the membrane. Next, in order to construct the trimeric assembly of the TM domain, multiple trimeric configurations of the TM helix were generated using Multimer Docking in ClusPro,^55^ a fast Fourier transformbased, rigid-body docking method. Briefly, a total of 10^9^ docking models were generated out of which a subset of 10^3^ models were selected based on the predicted binding-energy score of Cluspro. We then clustered these models using RMSD and ranked them according to their populations. The models in the top 10 clusters were visually inspected for the expected symmetry between the individual TM monomers and their membrane orientations, and the two best trimeric TM configurations were selected for additional refinement in MD simulations. The two selected trimeric TM configuation were similarly embedded in the membrane, solvated, energy-minimized, and equilibrated for 200 ns each. The stability of these models was evaluated by calculating the following metrics: i) RMSD of the TM domain with respect to the starting structure using *C_α_* atoms, ii) TM tilt/inclination, which was measured as the angle between the third principal axis of the moment of inertia of the heavy atoms in each TM monomer and that of the whole TM trimer, and iii) coordination number (coordNum implemented in the collective variables (COLVARS) module^56, 57^ of VMD^58^) quantifying the number of heavy atom pairs within a cutoff distance, between each TM monomer and the other two TM monomers, using a half-contact value of *d*_0_=4.5 Å. Here, for *d « d*_0_ the contact value is close to 1, at *d* = *d*_0_ the value is 0.5, and at *d » d*_0_ the value *∼*0. The total contact is then calculated by summing over all heavy-atom pairs in the two groups. The most stable TM trimeric conformation thus identified using the above three structural metrics was then used to construct the full-length spike structure.

### Constructing the C-terminal region

The C-terminal region, located in the lumen of the viral envelope, is predicted to be intrinsically disordered, through secondary structure predictions by JPred4, and does not have a suitable template structure. We used Robetta to generate multiple ab intio models of this region. From these, a model was selected based on its spatial arrangement with respect to the TM trimeric domain, namely, a model without any steric clashes when connected to the membrane-inserted TM trimer.

### Assembling the full-length spike

The structure of the full-length spike was constructed by assembling together the spike head and the individually constructed domains of the stalk region. The structures of the individual domains were aligned using VMD, with the sequential gaps between them ranging between 3-5 residues, which were filled using loop modeling in MODELLER.^59^

### Spike glycosylation and palmitoylations

The surface of the SARS-CoV-2 spike is known to be extensively glycosylated,^37^ but structural information on this feature is by and large missing from the available structures. Recent mass spectrometry studies have successfully identified both the spike’s glycosylation sites as well as their specific chemical compositions.^36, 37^ Utilizing this information, we added a total of 22 N-glycans and 1 O-glycan to each of the spike monomers (Table S2).

SARS-CoV-2 spike is also known to be palmitoylated at the endodomain,^35^ but its specific palmitoylation residues have not been determined. Since the cysteine-rich endodomain of the the SARS-CoV spike shares *∼*95% sequence identity with the SARS-CoV-2 spike, we used the palmitoylation data from the mutagenesis analysis of the cysteine clusters in the former^60^ as a reference for palmoytilation of the SARS-CoV-2 spike. Thus, we adopted 3 palmitoylation sites (residues 1,236, 1,240 and 1,241) in the first two cysteine-rich clusters of the C-terminal region in each spike monomer and covalently linked palmitoyl groups to them.

### MD simulations of membrane-bound spike

In order to probe the conformational dynamics of the entire spike as well as the motions displayed by the individual domains, we carried out MD simulations of the full-length, membrane-embedded spike structure, both in glycosylated and non-glycosylated forms. The following sections provide more details on the simulation system construction and the MD protocol used.

### System preparation

The final membrane-embedded, full-length spike system was prepared with the CHARMM-GUI webserver.^52, 53^ 22 N-glycosylations and 1 O-glycosylation were added to each spike monomer using Glycan Reader & Modeler in CHARMM-GUI.^28, 61^ Three palmitic acid tails were also added to cysteine residues 1,236, 1,240 and 1,241 in the endodomain of each monomer. A total of 15 disulfide bonds were introduced between specific cysteine pairs (15-136, 131-166, 291-301, 336-361, 379-432, 391-525, 480-488, 538-590, 617-649, 662-671, 738-760, 743-749, 840-851, 1,032-1,043 and 1,082-1,126). The protonation states of titratable residues were set using PROPKA.^62, 63^ Following this, the spike protein was inserted into a lipid bilayer with ERGIC composition described above.^22, 54^ The system was then solvated with water and ionized with 0.15 M NaCl, resulting in a simulation system with approximate dimensions of 250 *×* 250 *×* 390 Å^3^ with *∼* 2.3 million atoms.

### MD simulations

MD simulations were performed using NAMD2, a highly scalable MD engine.^64, 65^ The CHARMM36m force field was used to represent the proteins, lipids, glycosylations, palmitoylations, and ions,^66, 67^ and the TIP3P model was used for water.^68^ The simulations were performed as an NPT ensemble with the temperature and pressure maintained at 310 K and 1 bar using Langevin thermostat and barostat,^69, 70^ respectively. The SHAKE algorithm was used to constrain all bonds with hydrogen atoms.^71^ For the calculation of van der Waals interactions, a pairlist distance of 13.5 Å, a switching distance of 12 Å, and a cuoff of 10 Å were used. The Particle mesh Ewald (PME) method under periodic boundary conditions was utilized for the calculation of electrostatic interactions and forces.^72^

The system was equilibrated in 6 steps. In the first step, protein backbone and side chain heavy atoms were restrained using harmonic potentials with force constants of 10 and 5 kcal/mol/Å^2^, respectively, while the system was minimized for 10,000 timesteps using the steepest descent algorithm, followed by 100 ps of MD simulation. The modeled loop regions in the spike head and the linker regions connecting the different domain in the stalk were excluded from the restraints in this step to allow their adjustment. The restraints on the protein were sequentially reduced to half in each subsequent equilibration step, and the system was further equilibrated for 2 ns in each step. Additionally, coordinates of the lipid head group heavy atoms were restrained along the *z* axis (membrane normal) in order to maintain the membrane thickness during the whole equilibration phase with decreasing force constants from an initial value of 5 to 0.1 kcal/mol/Å^2^ over the 6 equilibration steps. The first 3 equilibration steps were carried out using a timestep of 1 fs and the subsequent steps and the production runs with a timestep of 2 fs. After the equilibration phase, all the restraints were removed, and a production run was carried out for 5 *µ*s.

As a control, full-length spike without glycosylations and palmitoylations was similarly prepared, embedded in an ERGIC-like lipid bilayer, equilibrated, and then simulated for 5 *µ*s.

### Analysis

We first evaluated the structural stability of the spike over the course of the MD simulations, specially in the modeled TM domain. The global conformational dynamics of the spike protein obtained from the simulations were characterized in terms of the bending and twisting motions of the spike head with respect to the different domains in the stalk region. Additionally, the potential role of glycosylation in modulating these motions was analyzed in terms of direct interactions with the lipid bilayer as well as contacts formed between the glycans at the interfaces of different domains and the lipids. Finally, we also investigated the potential effect of palmoytilations on SARS-CoV-2 spike in modulating the membrane curvature.

### Stability of the spike domains

The internal RMSD of different spike domains was calculated by superimposing the trajectory with respect to the starting structure of the respective domain using *C_α_* atoms. Furthermore, we characterized the stability of the modeled TM domain by calculating the following: i) TM tilt (described earlier for the evaluation of the TM domain models). ii) TM self-rotation which is calculated as the angle between the first principal axis of the moment of inertia of the heavy atoms in each TM monomer with respect to its equilibrated configuration, and iii) residue contact map between each TM monomer and the other two, calculated for C*α* atoms and using a half-contact distance of 4 Å.

### Global structural dynamics

The global conformational dynamics of the spike protein were characterized in terms of the orientational changes in the spike head with respect to the different stalk domains. For this, the trajectory was separately aligned using either the spike neck, the HR2 domain, or the TM domain, and the motion of the spike head was quantified with respect to the superimposed region in terms of the i) head bend, i.e., bending motion of the spike head, ii) head twist, i.e., twisting motion of the spike head in the *xy* plane, and iii) head distance to the superimposed region. For the calculation of the angles, specific vector representations were introduced for different domains as described below (see Fig. S1). For the calculation of the head bend, the spike head was represented by a vector connecting the *C_α_* atoms of the bottom and top residues of the central helices (i.e., residues 986 to 1,034), and the angle was calculated with respect to its initial position. For the calculation of the head twist, the top of the spike head was first approximated by a triangle with the *C_α_* atoms of residues 146 in the 3 spike monomers as vertices. The rotation of this triangle (quantified as the angle of one of its sides with regard to the initial position) was used as the twist angle. The head distance was calculated between the centroid of the triangle described above and the *C_α_* atom of the bottom residues of the different stalk domains used for superimposition (residue 1159 for the spike neck, residue 1204 for the HR2 domain, and residue 1236 for the TM domain).

### Multiple sequence alignment of human coronaviruses

In order to analyze the sequence conservation of the individual spike domains, multiple sequence alignment was performed for the protein from different human coronaviruses (HCoVs).

Currently a total of seven known HCoVs exist, namely, 229E and NL63 from the alpha sub-family of coronaviruses, and OC43, HKU1, MERS-CoV, SARS-CoV, and SARS-CoV-2, from the beta subfamily. Additionally, the bat coronavirus RaTG13, a closely related homolog of SARS-CoV-2, was also included in the sequence alignment. Multiple sequence alignment for the above eight sequences was carried out using the MAFFT program with the L-INS-i method^73^ and visualized using Jalview. ^74^

### Glycan-lipid and glycan-glycan interactions

As the glycan molecules attached to the HR2 domain are situated close to the membrane and to the glycans on the neck region, their possible interactions were further analyzed. For this, we identified all heavy atom contacts between glycans and other glycans or lipids (cutoff of 4.5 Å). The number of glycan heavy-atom contacts in each monomer for each frame in the trajectory was denoted as the glycan contact number, calculated separately for glycan-glycan and glycan-lipid interactions. Additionally, the bending of the HR2 domain with respect to the membrane as well as the angle between the neck region and the HR2 domain was also quantified. For the former, the trajectory was first aligned by rotating the first principle axes of the membrane patch to that in the first frame so that the membrane normal vector lied in the direction of the *z*. The HR2 bending was then calculated as the angle between the third principal axis of the HR2 domain, determined using the *C_α_* atoms, and the *z* axis representing the membrane normal. The neck-HR2 bending was determined similarly by obtaining the angle between the third principal axis of the HR2 domain and the neck region.

### Correlated motions in the spike

In order to identify potential correlations in the motions of the different domains of the spike, the Gram-Schmidt orthogonalization process was carried out, a widely used mathematical method to orthonormalize a set of vectors, in this case vectors representing orientations of the spike head, the HR2 domain, and the TM domain, generating 3 linearly independent, perpendicular vectors. For given vectors *..v*_1_ (HR2), *..v*_2_ (TM) and *..v*_3_ (spike head), representing the *C_α_* atoms of the HR2, TM and the central helices of the spike head, respectively, we define a plane formed by two vectors, e.g., *..v*_1_ and *..v*_2_, and obtain the components of *..v*_3_ on this plane and a perpendicular plane by performing orthogonalization. Thus, the spike head’s coplanar and orthogonal motions with respect to the HR2-TM domain bending can be derived to investigate the inter-correlation between different modes of bending.

### Calculation of membrane curvature

Since the palmitoyl tails have been shown to contribute to the bending of the membrane in other viruses,^32^ the curvature of the membrane around the fully-palmitoylated spike system was examined. Briefly, the coordinates of the upper and lower leaflet headgroups were first with a trigonometric surface function by the least square method, to maintain the periodic condition, followed by the numerical calculation of the mean curvature for the fitted grid points. For comparison, the membrane curvature of the non-palmitoylated spike system was also reported.

## Results and Discussion

SARS-CoV-2 binds to its host cell by the spike protein functioning as an extended viral antennae that recognizes the host cell surface receptors. Constructing the full spike and studying its behavior in its physiological environment remains the first and critical step in understanding the working of this most important component of the virus infection machinery. In the presented study, we first develop the full spike structure in its physiological, membrane-bound state, and subsequently, using *µ*s-scale MD simulations we characterize the conformational dynamics of the protein. We describe these conformational changes in terms of bending and twisting motions of the spike head with respect to the stalk, and examine possible correlations among inter-domain motions as well as the the potential role of glycan-lipid and glycan-glycan interactions in these motions. Sequence comparison with other human coronaviruses allows us to delineate multiple conserved hinge regions in the stalk domain that likely promote the conformational motions of the spike head. Furthermore, we quantify the effects of palmitoylation in the TM domain offering support for the role of this post-transnational modification in modulating the local membrane curvature.

### Construction of the membrane-embedded full spike protein

We combined homology modeling utilizing multiple template structures from homologous coronaviruses like SARS-CoV and MERS-CoV with fragment-based, ab initio modeling and secondary structure prediction to construct the missing regions in the spike head and the stalk including the HR2 domain (Fig. 1A and B and Fig. S2). Additionally, we carried out an expansive procedure for deriving a stable model of the TM domain, where we combine sequence-based secondary structure prediction for the TM region (Fig. S3), with protein-protein docking for generating the initial configurations of TM trimers (Table S3) followed by MD relaxation of suitable TM trimeric configurations in membrane (Fig. S4 and S5). The above-described modeling procedure was combined with the the biochemical data on the palmitoylation sites^35^ and the recently reported glycomics data to determine the glycosylation compositions of specific Asn residues^36, 37^ (Fig. 1C and Table S2), to develop the full-length, palmitoylated and fully-glycosylated spike structure (Fig. 1D).

We have also carried out structural comparison of our full-length spike model with previously published models.^28, 29, 75^ The spike head structure in our study is obtained from cryo-EM data as done in the previous models. The individual monomers of the spike head are observed to display relatively stable conformations during our simulations after their initial equilibration (Fig. S6). The HR2 domain in the previous studies was modeled using either a coiled-coil template from salmonella autotransporter adhesin,^29^ using a de novo coiled-coil modeling program,^75^ or using the HR2 template from SARS-CoV.^28^ Here, we also make use of the HR2 template structure from SARS-CoV,^47^ resulting in a model observed to be stable throughout the trajectory (Fig. S6). As SARS-CoV and SARS-CoV-2 display complete sequence identity in the HR2 domain, our stable model is expected to provide a closer physiological representation of the HR2 domain compared to the coiled-coil models used in the earlier studies. The TM domain in the previous spike models^28, 29^ was constructed using a template structure from HIV.^50^ It is noteworthy that the TM helices in HIV each contain a ‘GxxxG’ motif at their N-terminal end, a motif known to be frequently involved in close inter-TM helix interactions.^76^ Consequently, the TM domain assembly in HIV displays inter-helical contacts in this region with the C-terminal half only held together by polar contacts.^50^ As the ‘GxxxG’ motif is present in the central region of the TM helices in the SARS-CoV-2 spike (Fig. 1A), the contacts between the TM helices in the SARS-CoV-2 spike can be expected to lie in this region instead. The TM domain in our spike model is a trimeric assembly with the majority of the inter-helical contacts formed between the central regions of the helices containing the ‘GxxxG’ motifs (Fig. 2A and B). Additionally, the TM domain maintains a relatively stable and symmetric structure during the simulations (Fig. S6 and Fig. 2C, D, E and F). The correct prediction of the different stalk domain structures, including the trimeric HR2 and TM domains, may be critical to correctly describing the global motions observed in the spike head, as described below.

**Figure 2:**
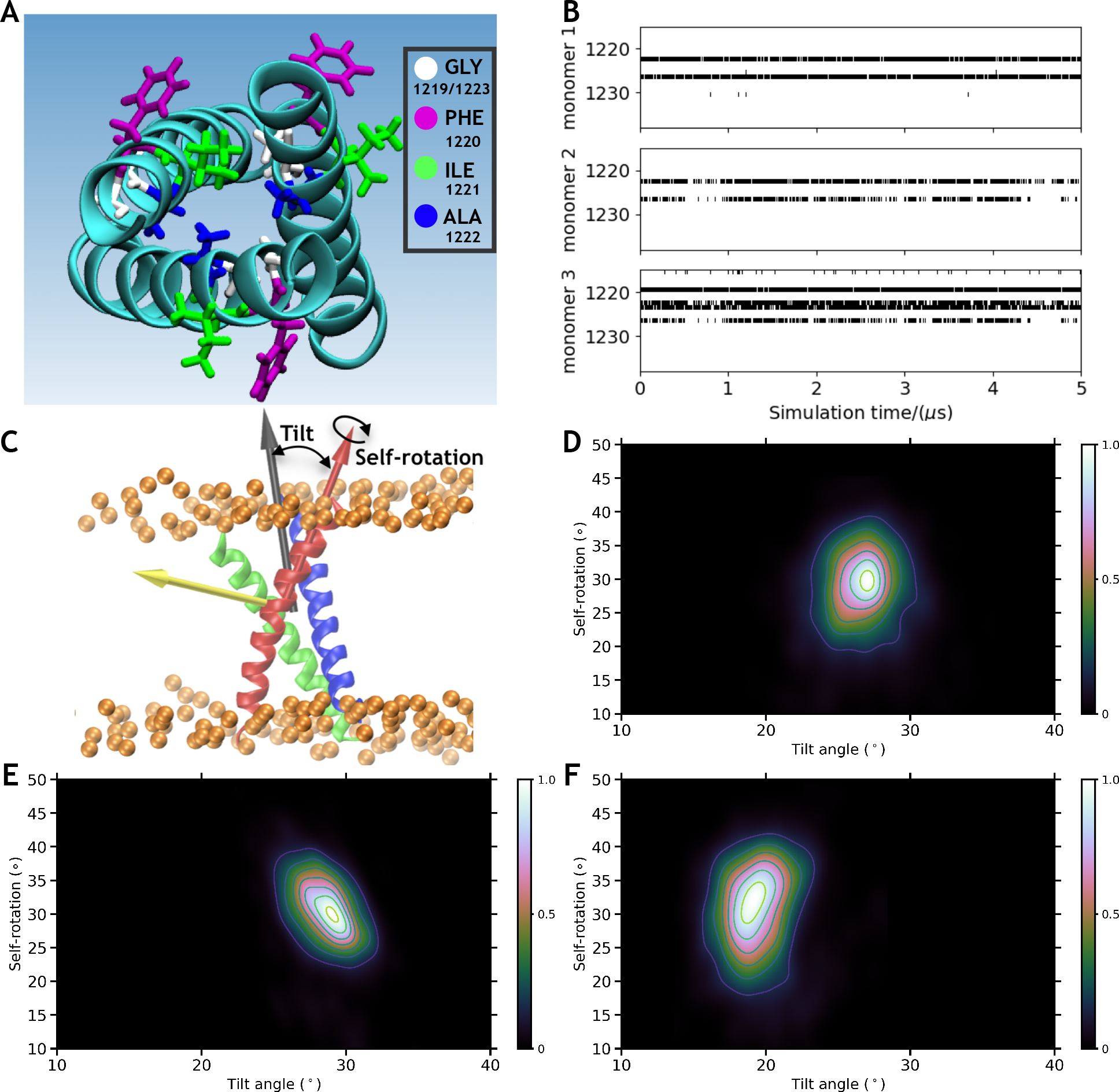
Stability of the modeled TM domain: (A) The top view of the modeled TM domain as well as the licorice representation of the ”GxxxG” motif (residue 1,219-1,223) are shown. (B) Contact maps between a TM monomer and the other two monomers in the TM domain during the trajectory. A black dot is used to indicate the presence of at least one *C_α_*-*C_α_* contact (4.0 Å cut-off) between residues of a monomer and the other two monomers. (C) TM monomers’ self-rotation and tilt motions. The self-rotation of each monomer is calculated with respect to its equilibrated configuration, and the tilt angle is measured with respect to the whole TM domain. The heat map for the distribution of the tilt angle versus the self-rotation angle is shown for the TM monomer 1 (D), monomer 2 (E), and monomer 3 (F).

### Bending and twisting motions of spike head around stalk

To further describe the the dynamics employed by the spike to sample the crowded cellular surface, we characterized its global motions in terms of the movements of the spike head with respect to the stalk region during the simulations (see Supplementary Video 1). These motions were quantified in terms of head-twist and head-bend angles as well as head-distances calculated separately with respect to either the spike neck, the HR2 domain, or the TM domain (Fig. 3A). In its fully-glycosylated, native form, the spike can sample large head-twist angles (in the *xy* plane) with respect to both the HR2 and TM domains, highlighting the large conformational flexibility available along this degree of freedom (Fig. 3B top, F and G). A wide normal distribution of head-twist angles is observed with respect to the HR2 domain, with the highest sampled configurations turned by almost 90*^◦^* relative to the starting structure. With respect to the TM domain, although lowly populated, spike-head can sample conformations that are almost oppositely facing (turned by as much as 160*^◦^*) compared to the starting orientation. A relatively smaller range of motion is allowed around the spike neck, with the most populous regions lying close to the starting structure (Fig. 3B top and E). These results can be directly related to the length of the linkers connecting the different domains of spike. The domains around which spike head displays larger motions (HR2 and TM domains) are connected by longer linkers (linker 2: 13 residues; linker 3: 11 residues), compared to the domain around which spike head displays relatively more restricted motions (spike neck), that is connected to the spike head by a smaller linker (linker 1: 7 residues). Furthermore, direct comparison of the head-twist motions in the glycosylated form to the control non-glycosylated system shows surprisingly that these motions become restricted in the latter (Fig. 3B bottom), especially with respect to the head-twist angles calculated around the HR2 and TM domains. These results point to an unexpected enhancing role of glycans in the overall dynamics of the spike.

**Figure 3:**
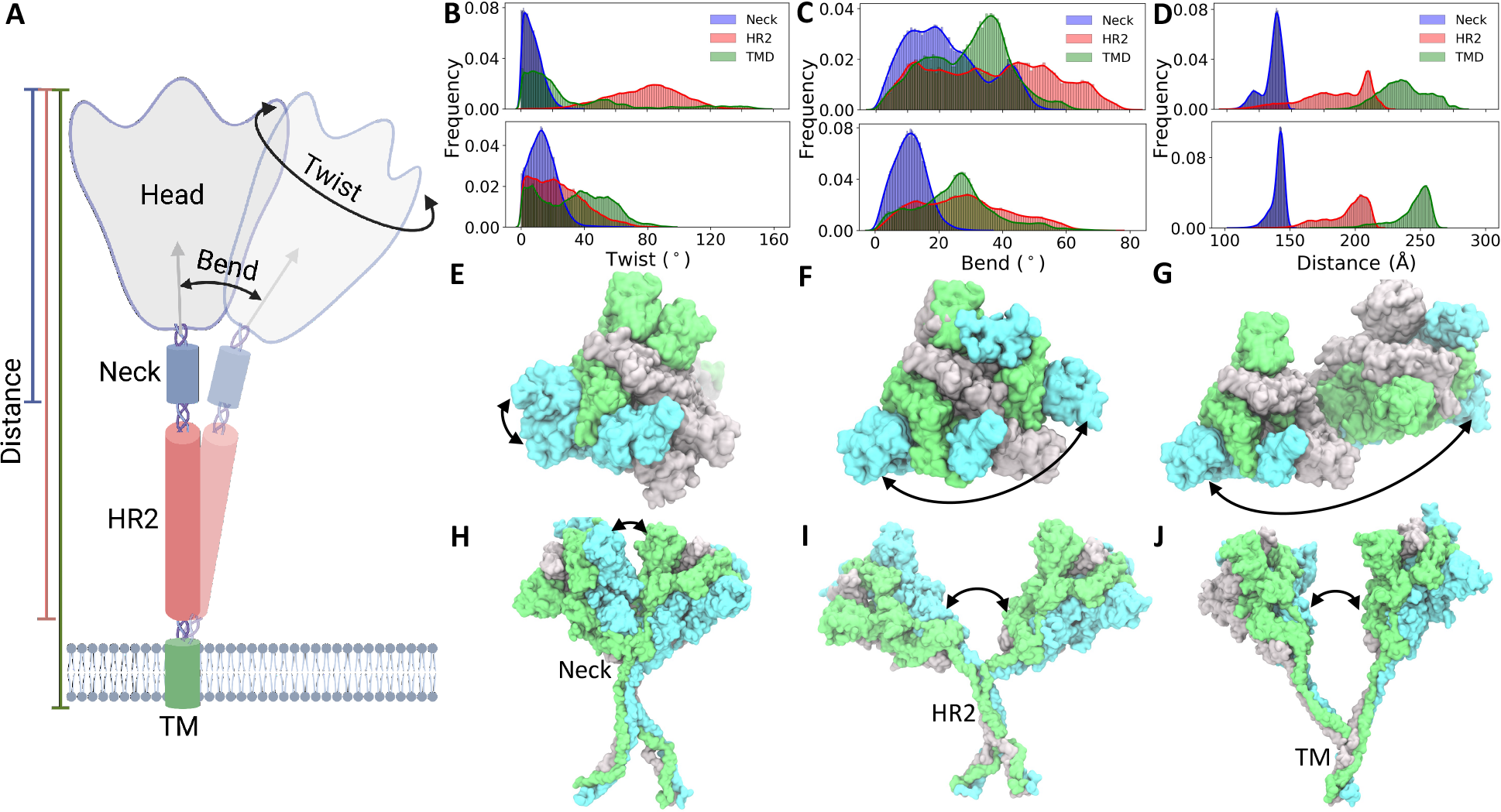
Global conformational dynamics of spike: (A) The schematic depicts the calculated bend and twist motions of the spike head (grey) around the spike neck (blue), the HR2 domain (red) and the TM domain (green), as well as the calculated head distances with respect to these domains. The probability distribution (in terms of normalized frequency) of the (B) twist, (C) bend, and (D) distance of the spike head with respect to the spike neck, HR2 domain and TM domain are provided. The top and bottom plots show the distributions observed in the fully-glycosylated spike and the non-glycosylated spike. Example snapshots of the spike (surface representation with each monomer in a different color) displaying the largest head twist with respect to the (E) spike neck, (F) the HR2 domain, and (G) the TM domain are shown for the fully-glycosylated system. Similarly, example snapshots of the spike showing the largest head bend with respect to the (H) spike neck, (I) the HR2 domain, and (J) TM domain are shown.

Similarly, the spike head displays a broad range of head-bend angles with respect to the HR2 domain, sampling configuraion almost orthogonal to the starting structure (Fig. 3C top and I). Comparatively, smaller head-bend motions are observed around the spike neck and the TM domain, with the respective populations showing distributions over a range of 0 *−* 60*^◦^* angles (Fig. 3C top, H and J). The extent of these bending angles is also directly related to the changes in the head-distances with respect to these domains, with the largest distance changes observed in conformations with the largest head-bend angles (Fig. 3D top). Thus, the spike head can move as much as 125 Å with respect to the HR2 domain compared to its starting configuration. Again, comparison of the glycosylated and non-glycosylated forms shows that the spike domains rigidify (display relatively smaller range of motion) in the absence of glycans (Fig. 3C and D bottom). The potential role of the glycans in modulating global motions of the spike is further discussed in later sections.

Mechanistically, these motions (bend/twist/translation) of the spike head represent major degrees of freedom available bestowed upon the receptor-binding domain (located in the spike head) to locate and bind to the target ACE2 receptors in the host cell membranes. As spike proteins have been found to be sparsely distributed on the virion envelope (recent cryo-EM and tomography studies^12, 77^), the flexibility and the allowed range of motions of the spike domain may play an important role in extending the search space covered by the spikes. Cryo-EM and other computational studies have similarly reported the conformational diversity of the spike head around its stalk.^75, 78^ Compared to our results, the conformational variability of the spike head was observed to a smaller extent in one of these MD studies,^75^ likely due to the crowded conditions used in the simulation system, where 4 spikes were packed at close proximity to each other, preventing the protein to amply sample the available space. The crowded distribution used in the above study is unlikely to represent the physiological distribution observed in the whole virion cryo-EM study where approximately one spike is observed per 1000 nm^2^ of the viral surface area.^77^ Additionally, previous studies have quantified the inter-domain motions compared to the spike head orientational changes measured in our study, that are more directly related to the motions of the RBD binding to the host receptors, and hence directly related to the function of the protein.

### Sequence conservation in multiple hinge regions

In order to examine potential generality of the observed global motions in the SARS-CoV-2 spike, we analyzed the sequence similarity of the spike protein, specifically focusing on the flexible regions connecting the different domains. These include the small connecting linkers between the spike head and the neck (linker 1: NTVYDPL), between the spike neck and the HR2 domain (linker 2: HTSPDVDLGDISG), and between the HR2 domain and the

TM domain (linker 3: GKYEQYIKWP). The three linkers are found to be rich in Pro and Gly, residues known to contribute to the conformational fly in proteins^79, 80^ (Fig. 4A). Additionally, the Pro residues in these regions may be important for the hinge or recoil motion suggested to take place in the refolding of the spike during the fusion process, which may result in an elongated post-fusion state of the protein.^15^

**Figure 4:**
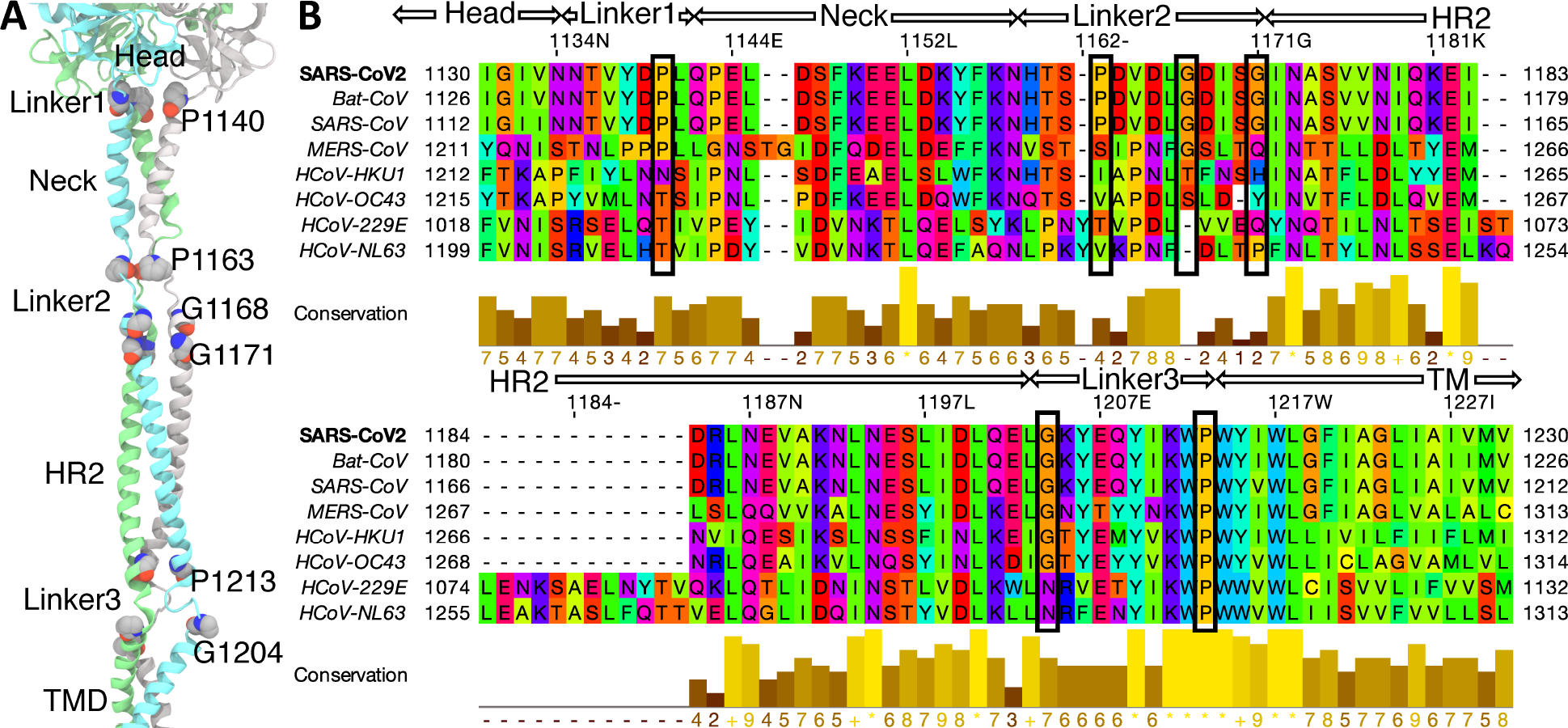
Multiple sequence alignment of spike homologs. (A) Structure of the stalk region of spike in our model is shown. Pro and Gly residues of the linkers are shown in van der Waals representation. (B) The sequence of SARS-CoV-2 spike is aligned with those from closely related virues (bat coronavirus RaTG13 and six other human HCoV viruses). The residues are colored according to their physiochemical properties in different shades of rainbow colors (Taylor coloring scheme). The different domains and linkers are labeled at the top of the sequence. The completely and partially conserved Pro and Gly residues present in the flexible linkers are enclosed in black boxes. Conservation between the different sequences is visualized as histograms with their heights, and color variation from brown to yellow, reflecting the level of conservation of physicochemical properties. Conserved and identical amino acid columns show the highest score of 11 and are indicated by ’*’, whereas conserved and similar amino acid columns show a score of 10 and are indicated by ’+’. Other groupings are indicated by lower scores accordingly.

To further elucidate the possible functional importance of these residues in the mobility of the spike, multiple sequence alignment was carried out with other coronaviruses infecting humans. Interestingly, all Pro and Gly residues present in the flexible regions are completely conserved in the homologous Bat-CoV and human SARS-CoV, and to a large extent in human MERS-CoV where only Pro-1163 is not conserved (Fig. 4B). Most of these residues are not conserved in other human coronaviruses except Pro-1213 that is completely conserved in all coronaviruses. We propose that the presence of these conserved flexible residues in the regions connecting the different domains not only promotes the observed conformational flexibility of the spike head around the stalk domains but, compared to other coronaviruses, also provides an evolutionary advantage to SARS-CoV-2, SARS-CoV and MERS-CoV, coronaviruses known to be highly infective in humans.

### Stalk bending motions modulated by glycan-lipid and glycan-glycan interactions

In addition to the important roles glycans can play in shielding the spike from neutralizing antibodies,^21^ we noted that the glycans also form contacts with each other, as well as with the lipid bilayer, during the simulations (see example of these phenomena in Supplementary Video 2). To obtain deeper insights into the impact of these glycan interactions, especially with respect to the spike global motions, we evaluated the relationship between the inclination of the different stalk domains and the glycan-lipid or glycan-glycan contacts formed, both in the native glycosylated spike simulations and the control non-glycosylated spike. In the glycosylated spike when the inclination angle of the HR2 domain exceeded a threshold of around 20*^◦^*, we observed formation of more glycan-lipid or glycan-glycan contacts, indicating a correlation between the bending of the HR2 domain and the glycan interactions (Fig. 5A, B, C and D). On examining the distribution of the HR2 tilt angles towards the membrane in the glycosylated spike simulation, only *∼*20% of the trajectory was found to lie between 0*^◦^* – 20*^◦^* (vertical orientation) compared to *∼*80% of the non-glycosylated trajectory that lied below the 20*^◦^* inclination. It can be inferred that in the fully-glycosylated spike when no glycan-lipid interactions are formed the HR2 domain can move freely within the 20*^◦^* inclination (Fig. 5A and B); the glycan-lipid interactions appear when the inclination angle lies between 20*^◦^* – 50*^◦^*, further restraining the flexibility of the HR2 hinge motion in this range. The HR2-neck inclination shows similar properties as the HR2-membrane inclination, where larger inclination angles, ranging between 20*^◦^* – 90*^◦^*, mostly correspond with high glycan-glycan contacts (Fig. 5C and D). Although the HR2 domain in the non-glycosylated spike can show a similar range of motion (with respect to the neck) as in the native form, almost half of the sampled configurations lie close to the starting vertical orientation (0*^◦^* – 20*^◦^*) (Fig. 5A and C), suggesting coupling between glycan-glycan interactions and the dynamics of the stalk. Overall, the data indicate that the glycans around the flexible linkers can further enhance the relative motion between the stalk domains, in turn increasing the probed space by the spike head.

**Figure 5:**
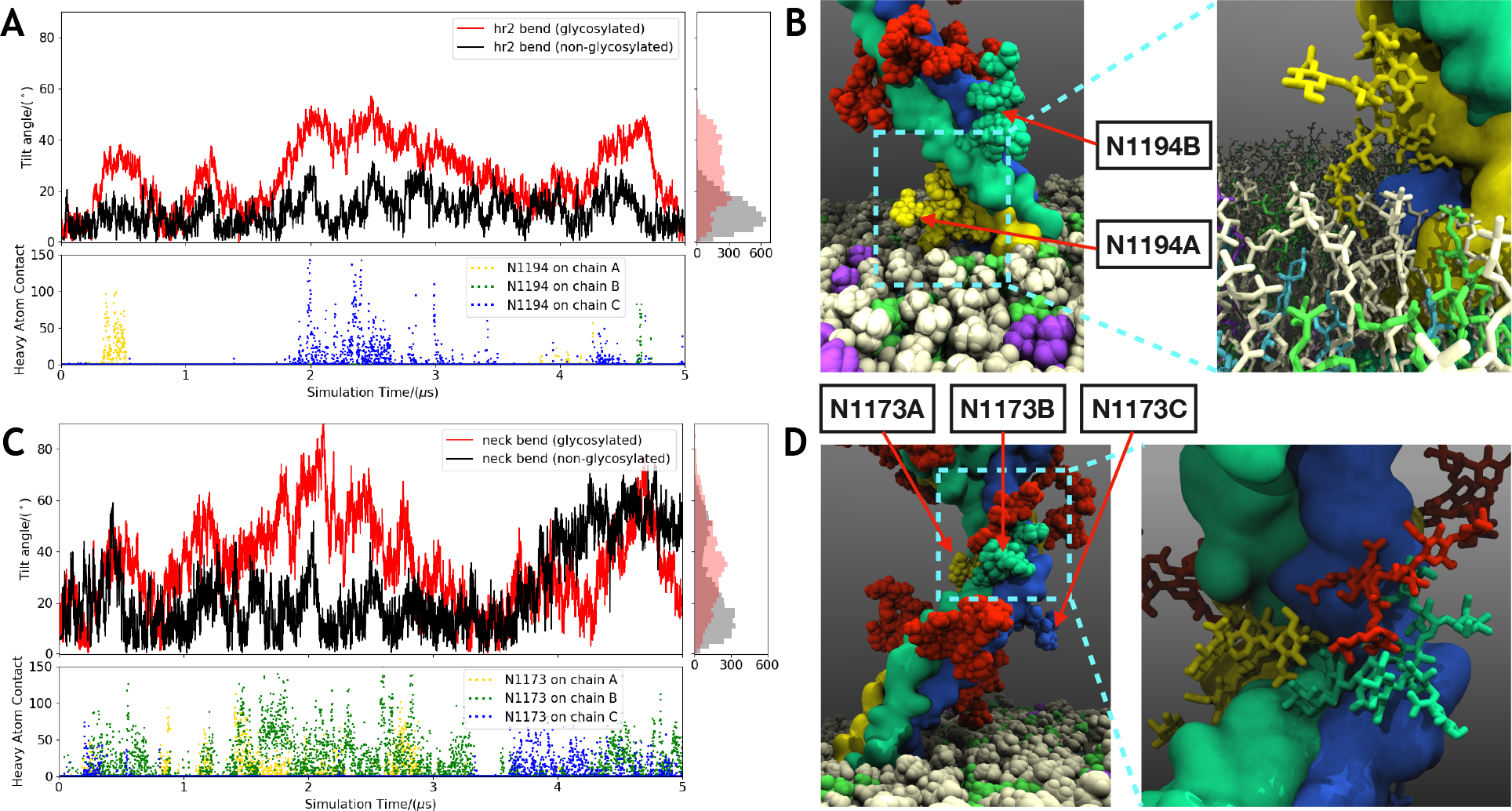
Glycan-glycan and glycan-lipid interactions: (A) Time series of the HR2 tilt angle towards the membrane in the glycosylated (red) and non-glycosylated (black) simulations (distribution histogram shown on the side). The number of heavy-atom contacts (within 4.5 Å) between the glycans at N1194 and lipids over the course of the trajectory is also shown (bottom). The contacts shown use the same color scheme as the corresponding monomer and glycan representations shown in (B) and (D). (B) Snapshot of the glycan at N1194 contacting with lipids in van der Waals representation and zoomed in detail shown in inset in licorice representation. Glycans in the front are labeled. (C) Similar tilt angle between the HR2 domain and the neck region of the glycosylated (red) and non-glycosylated (black) simulations along with the distribution histogram (side). The glycan-glycan heavy atom contacts between N1173 on HR2 and N1158 on the neck region is shown as points (bottom). (D) Snapshot of the glycan at N1173 (same color scheme as chains) contacting with glycans at N1168 (red) in van der Waals representation and zoomed in detail shown in inset in licorice representation.

N-glycans have been reported to be involved in mediating receptor recognition and facilitating cell adhesion.^81^ The self-recognition and conjugation has been previously demonstrated for N-linked complex glycans with multiple N-acytylglucosamine (GlcNAc) termini. ^82, 83^ The glycans located at the bottom end of the neck region (N1158), as well as the top (N1173) and bottom (N1194) ends of the HR2 domain, are all complex glycans containing multivalent GlcNAc (Table SS2). Additionally, these glycans are completely conserved in human coronaviruses (Fig. 4B). Overall, we propose that these complex glycosylation sites not only shield the stalk from epitopes, by occupying the space around the flexible hinges connecting the neck, HR2 and TM domains, but also contribute to the flexibility of the stalk, that may in turn increase the range of motion of the spike head allowing for more effective sampling of the human cell surface.

On further examination of the glycan-lipid interactions we observed that the majority of these contacts are formed with the zwitterionic lipids (POPC/POPE) followed by anionic lipids (POPS) (Fig. S7). Majority of the interactions between glycans and lipids consist of hydrogen bonds formed between O/N atoms of the N-acetyl-D-neurominic acid (NE5AC) or D-galactose sugars present in the glycans and N atoms from PE head group or O atoms in the PE/PC phosphate groups. These interactions may further stabilize the inclined conformations of the HR2 domain with respect to the membrane. The contacts with cholesterol are minimal, which is expected as cholesterol is known to insert more deeply in the membrane core compared to the phospholipids.^84, 85^

### Correlated inter-domain motions in the spike

The global conformational dynamics of the spike were further examined for the presence of correlated motions between its different domains. A correlation between the motions of the spike head and the TM domain about the HR2 domain is evident in the fully-glycosylated spike (Fig. 6A and B; see Supplementary Video 3). A high Pearson’s coefficient of 0.85 between the Head1 and TM vectors (Fig. 6A and B), representing the spike head’s coplanar motion with respect to the HR2-TM domain bending, further confirms the observed correlation in the motions of these domains. A Pearson’s coefficient of 0.48 was also observed between the Head2 and TM vectors (Fig. 6A and B), showing that the spike head’s orthogonal motion is similarly correlated, but to a lesser degree, with respect to the HR2-TM domain bending. This is an interesting observations in that the orientational motions of the different domains of the spike appear to remain inter-related even though they are connected by flexible hinge regions. Without glycosylations, not only is the magnitude of inter-domain motions dampened (Fig. S8), but, at the same time, the correlation between them is also reduced, with the non-glycosylated spike showing relatively lower Pearson’s coefficient, 0.78 and 0.39, respectively, for the in-plane and orthogonal motions of the spike head with respect to the HR2-TM domain bending. In general, the correlated motions between the different spike domains can be attributed to its unique tertiary structure and can be considered an intrinsic property of the protein, but, at the same time, glycosylation may further amplify these motions through the extensive glycan-lipid and glycan-glycan interactions in the system (Fig. 5). For example, interactions of the glycans located at the bottom end of the HR2 domain with the lipids seem to promote its bending towards the membrane that may in turn facilitate formation of more contacts between the glycans located at the top/opposite end of the HR2 domain and glycans in the neck region, promoting bending of the spike neck and head in the opposite direction.

**Figure 6:**
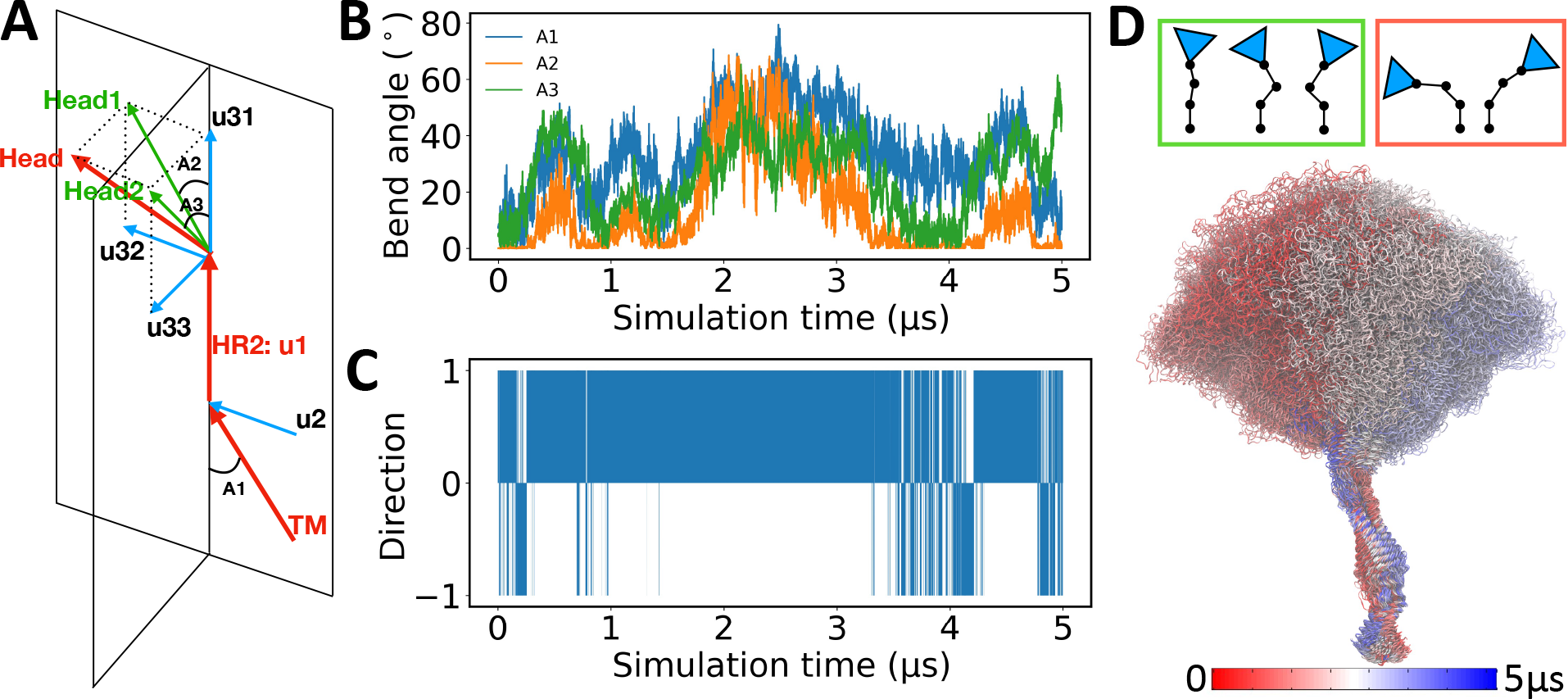
Global dynamics correlation in spike: (A) Vectors representing the different domains of the spike (arrows in red): spike head (top), HR2 (middle), TM (bottom), and their orthogonalized sub-vectors (arrows in blue) in orthogonal planes of spike head (u31, u32 and u33) and TM (u2). Head1 and Head2 (arrows in green) are respectively the portions of the head vector on the HR2-TM plane and the one perpendicular to it, representing the lateral and axial movements of the head. (B) The time-series of the A1, A2 and A3 angles introduced in (A) are shown for the fully-glycosylated spike. (C) The time-series of the Gram-Schmidt process for computing the dot product between the u2 and u32 vectors in (A) is shown. A value of +1 means the vectors are pointing in the same direction and a value of -1 means that they are pointing in the opposite direction. (D) The conformational ensemble of the spike generated by taking 100 images evenly distributed over the trajectory of the fully-glycosylated spike and superimposed on the stalk region (the HR2 and TM domains). The color changes from red to blue over the time. Cartoon representation of the spike on top (spike head depicted as triangle and different stalk domains connected by lines) in green box are the favored (highly sampled) conformations, whereas representations in red box are rarely sampled during the simulations.

The Gram-Schmidt analysis (see Methods) indicates concerted motions of the spike head and the TM domain about the HR2 domain. Here, the orthonormal vectors representing the spike head and the TM domain are oriented in the same direction about the HR2 domain for 88% of the simulation in the fully-glycosylated spike (Fig. 6C). Non-glycosylated spike displays a similar behavior, with the the orthonormal vectors oriented in the same direction for 89% of the time, further highlighting that these motions are largely a property of the spike structure itself. Topologically, this reflects a configuration where the spike head and the TM domain orient diametrically opposing each other around the HR2 domain, a configuration which may further extend the reach of the spike head and the RBD for the target cellular receptors (Fig. 6D). Additionally, considering the fact that the inclination of TM domain in the membrane can be substantial, ranging between 0-30*^◦^* (Fig. S9), the concerted, opposing bendings of the spike head and the TM domain may pull the head back to an upwards position with respect to the membrane, preventing it from severe inclination and falling into the membrane and thus maintaining a proper configuration for binding with the ACE2 receptor.

### Role of palmitoylations in modulating membrane curvature

Protein S-palmitoylation, the covalent lipid modification of the Cys residues, widely exists in viral membrane proteins and reported to contribute to membrane fusion and other processes.^31–33^ To characterize the effect of spike palmitoylations on the membrane shape, we compared the membrane curvature for the control simulation of non-palmitoylated spike with the palmoytilated (and fully glycosylated) one, taking into account the periodicity in fitting the lipid head groups (Fig. 7A, B). The two simulations share a similar membrane size, but the palmitoylated system seems to deform the membrane more than the non-palmitoylated system (Fig. 7C). Keeping in mind that the periodic boundary conditions used in the simulations can suppress long-range curvature effects, comparing the local curvature (in the range of 20-50 Å from the center of the TM domain) shows that the palmitoylated TM domain is found within a positively curved membrane, whereas the non-palmitoylated TM prefers a saddle point. Thus, the palmitic tails may induce a positive curvature locally in the vicinity of the TM domain.

**Figure 7:**
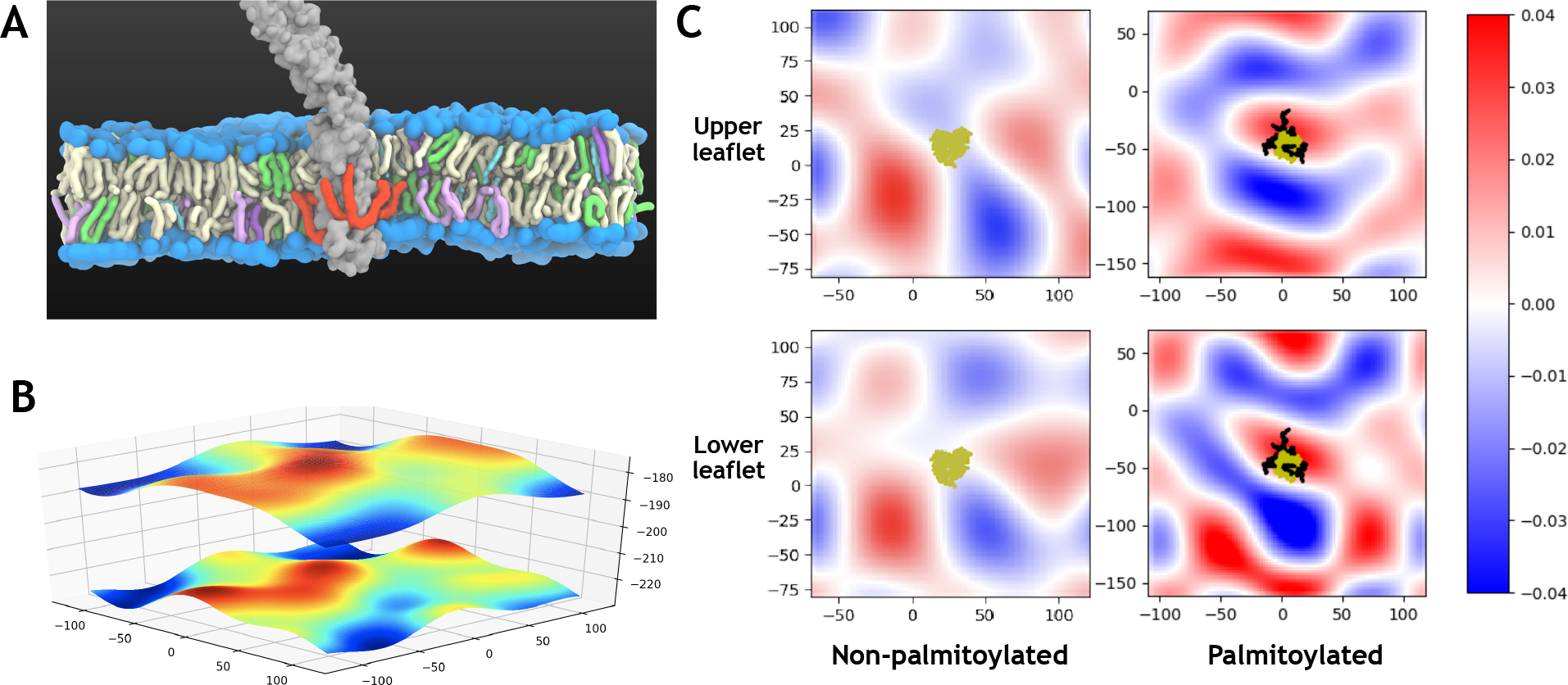
Membrane curvature modulation by palmitoylation: (A) Section view of the membrane embedded palmitoylated spike is shown. The coloring scheme for the lipid composition is identical to Fig. 1D, with the palmitoylations and lipid head groups displayed in red and blue surfaces, respectively. (B) Fitted surface (periodic) with lipid head group coordinates used for the calculation of curvature. (C) The mean curvature of the fitted membrane surface of the upper and lower leaflets as in (B). The left and right panels show the curvature profiles at the end of the simulations of non-palmoytilated and palmitoylated TM domains. The heavy atoms of the TM domain (yellow) and palmitoyls (black) are also shown.

The lipidation of other viral proteins is relatively well studied. In influenza virus, cryo-ET measurements have shown membrane curvature induced by HA palmitoylation.^32^ Apart from this, non-acylated HA mutants of influenza virus show a severe negative effect on the membrane fusion step of the infection process.^86^ As shown by the results above, SARS-CoV-2 spike palmitoylation may similarly contribute to the bending of the membrane, especially locally, and thus play a role in the viral fusion with the host cell membrane.

## Conclusions

The spike protein of SARS-CoV-2 constitutes a major component in the viral infection and, therefore, the main target for both diagnostics and vaccines developed for the disease caused by the virus. To better understand the many mechanistic steps the spike is involved in, it is imperative to characterize its structure and dynamics in the most detailed manner and under the most realistic conditions possible. We report here a full model for the full-length spike in its native, glycosylated and palmitoylated, membrane-bound form which we use for several microseconds of atomistic simulations highlighting some of the dynamics employed by the spike to search for and locate the host cell receptors effectively (Fig. 8). The hinge regions identified in the stalk domain directly modulate the conformational landscape of the spike head, where the RBDs reside, and may thus represent important structural elements regulating the effective role of the spike in binding its receptor. These highly conserved hinges may offer novel points of target for developing alternative drugs and antibodies that bind to and modulate the structure/dynamics of these regions, and therefore the effectiveness of the spike protein. In addition we propose possible roles for posttranslational modifications of spike, specifically glycosylation and palmitoylation, in modulating its dynamics. The results provide a deeper understanding of the intricate roles and functions of the different elements of the spike, each evolved in order to maximize the chance of successful infection and transmission of the genetic material to the host cell. Future extended simulations studying the spike behavior in a full viral envelope will allow further characterization of the impact of a more crowded environment of the envelope and its proteins in modulating the structure and dynamics of the spike and will provide additional insight into the inner working of the virus.

**Figure 8:**
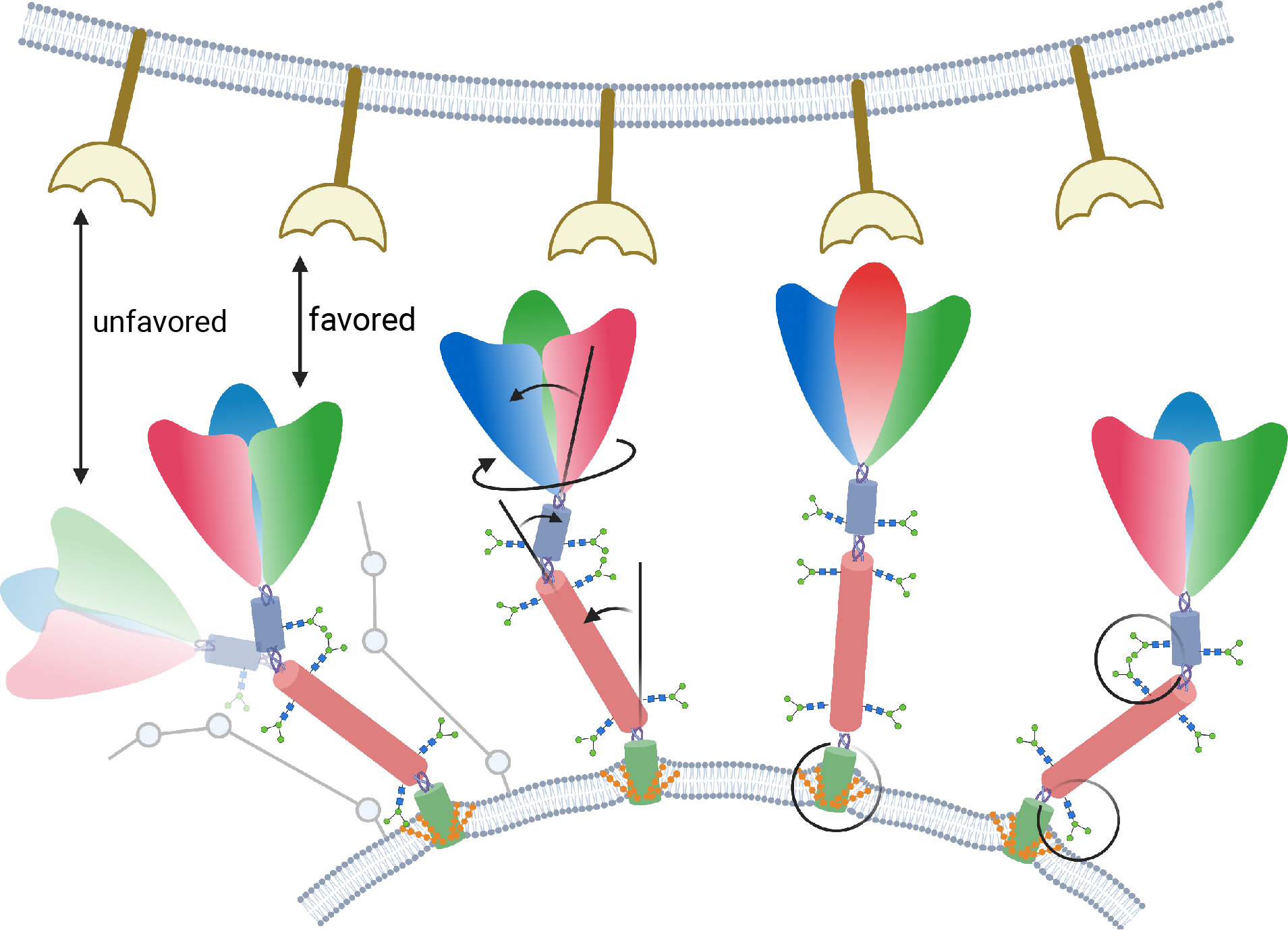
Impact of post-translational modifications on spike global dynamics and mem-brane deformation. The host cellular membrane with binding receptors are shown at the top, and the viral envelope containing multiple spikes at the bottom. Each spike head is shown in three colors, representing the three constituting monomers, and the different stalk domains are also shown in different colors. Spike copies shown from left to right highlight the following results: i) correlated motions of the spike head and stalk allowing reorientation of the spike head with respect to the receptor (stick representations of the spike in favorable and unfavorable orientations also shown on the side); ii) global motion of the spike head around the stalk (twisting and bending motions) allowing it to broaden its reach for the cellular receptors and optimally binding to it; iii) membrane curvature induced by palmitoylation potentially plays a role in viral fusion; iv) glycan-lipid and glycan-glycan interactions restricting/modulating the stalk motions. Created with BioRender.com.

## Acknowledgement

This study was supported by the National Institutes of Health under awards P41-GM104601 and R01-GM123455 (to ET). Simulations were performed using computational resources provided by Lawrence Livermore National Laboratory, Oak Ridge Leadership Computing Facility, and Microsoft Azure, under COVID-19 HPC Consortium Grant MCB200183 (to KK).

## Supporting Information

**Figure S1:**
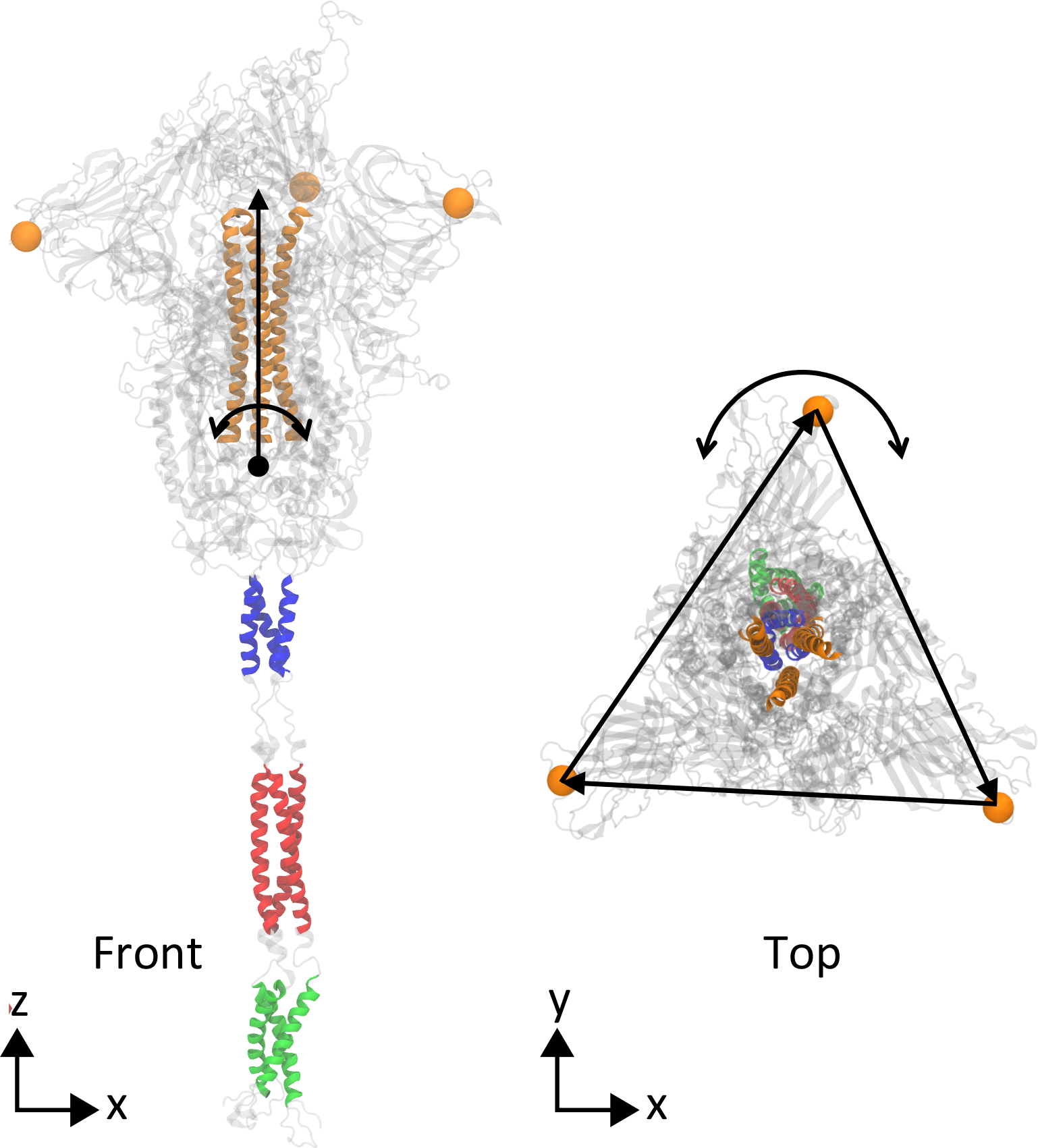
Selection for global analysis. Front and top views of describing selections used to represent the different domains in the calculation of respective vectors. TM: green, HR2: red, spike neck: blue, and spike head (represented by central helices): orange. For the calculation of the head twist, the top of the spike was approximated by a triangle formed by the C*_α_* atom of residue 146 from each monomer (orange van der Waals sphere). The sides of this triangle are the vectors used to calculate the bending and twist angles with respect to the starting geometry.

**Figure S2:**
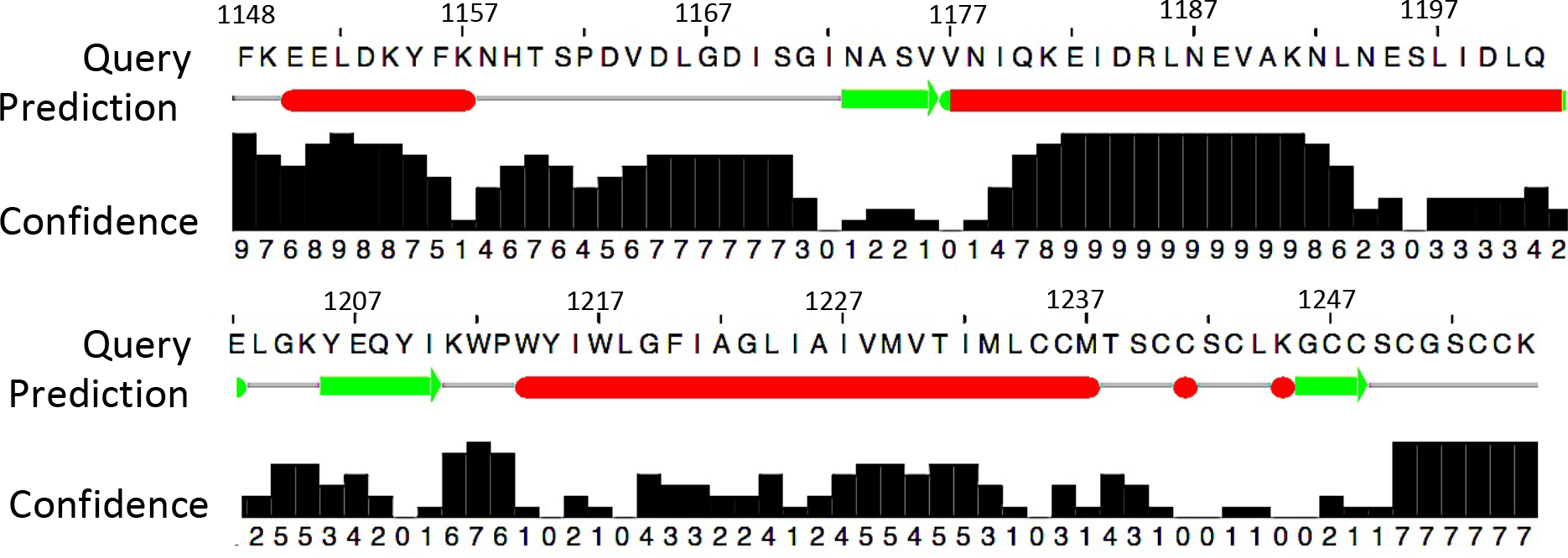
Secondary structure prediction for the unresolved stalk in spike. generated using JPred4. ^48^ Red bars represent predicted helical structures, and green bars represent *β−*sheets. The confidence level for the predictions ranges between 0 (low confidence) and 9 (high confidence).

**Figure S3:**
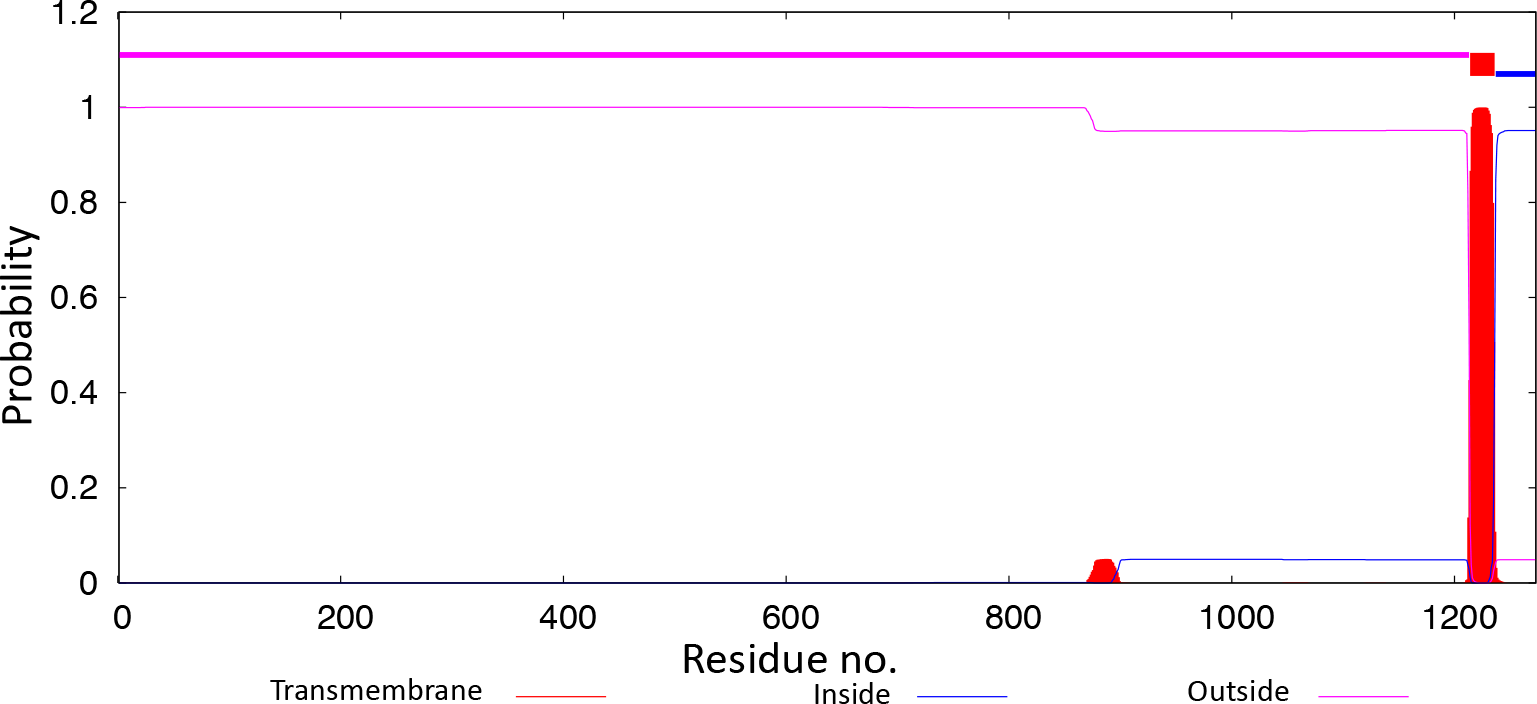
Spike TM domain prediction using TMHMM. ^51^. The plot shows the predicted probabilities of transmembrane as well as intracellular and extracellular regions of the protein sequence. At the top of the plot (between 1.0 and 1.2) the most probable location of the protein sequence (described in the bottom legend) is shown.

**Figure S4:**
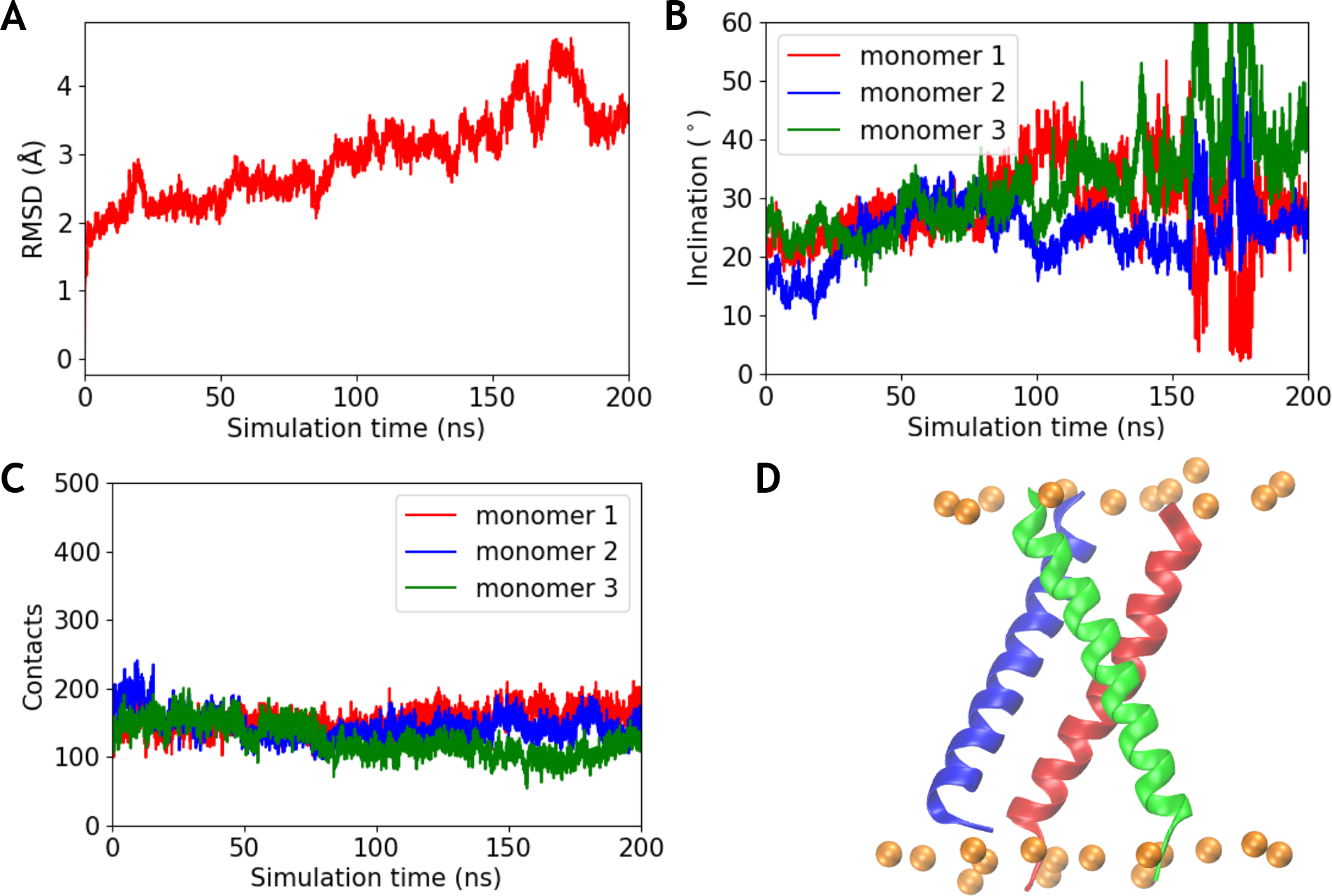
Stability of predicted TM domain model 1. (A) RMSD of the simulated TM domain model 1 (refer to Table S3), after aligning The trajectory with the starting structure using *C_α_* atoms. (B) The inclination of each TM monomer calculated with respect to the whole TM domain during the simulation. (C) The number of contacts between one TM monomer and the other two monomers in the TM trimer, quantified in terms of the coordination number (described in methods). (D) Snapshot of the TM domain structure at the end of the simulation is shown. The TM monomers are shown in same colors used in (B) and (C), and the phosphorus atoms of the lipid bilayer are shown as orange spheres.

**Figure S5:**
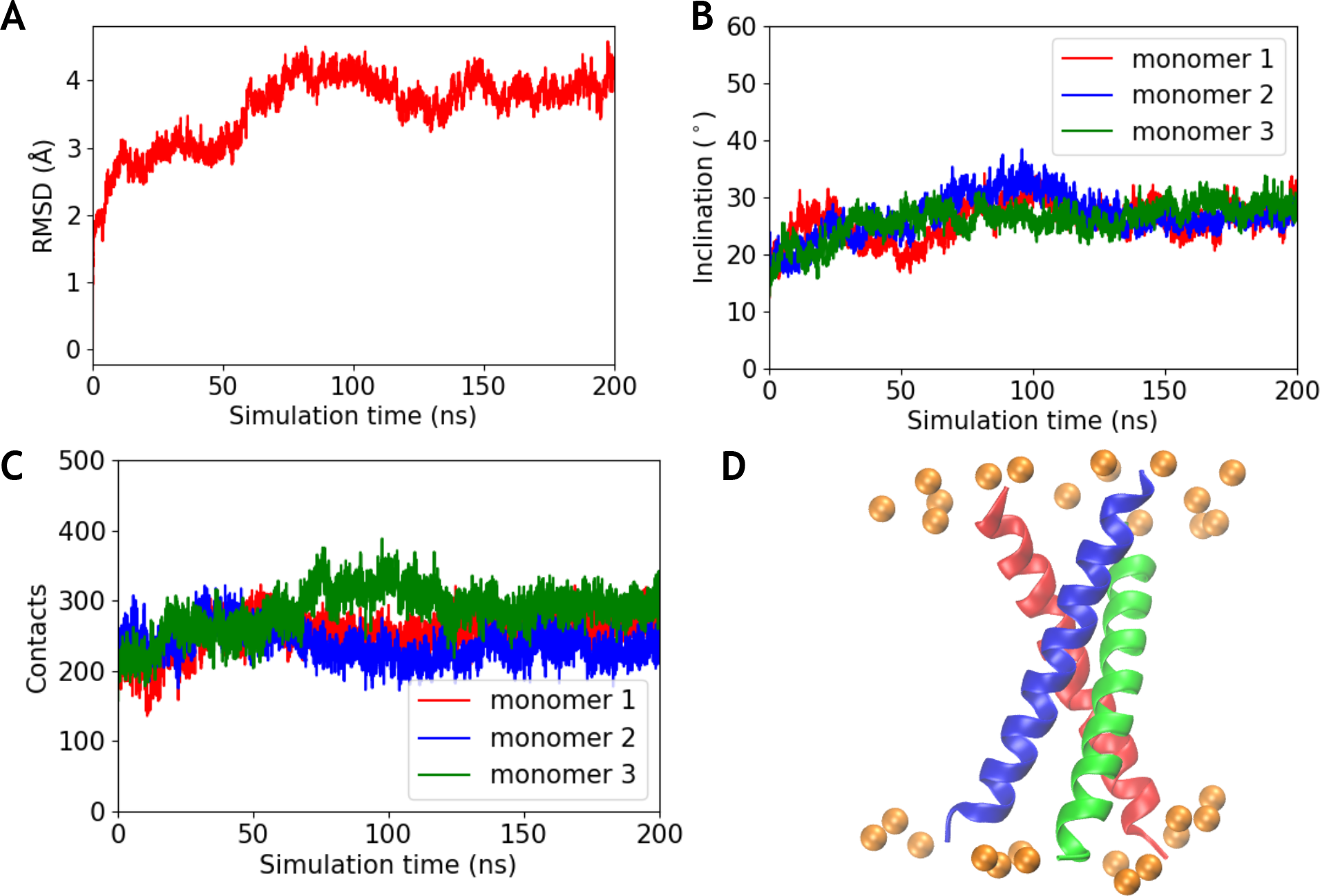
Stability of predicted TM domain model 3. (A) RMSD of the simulated TM domain model 3 (refer to Table S3), after aligning The trajectory with the starting structure using *C_α_* atoms. (B) The inclination of each TM monomer calculated with respect to the whole TM domain during the simulation. (C) The number of contacts between one TM monomer and the other two monomers in the TM trimer, quantified in terms of the coordination number (described in methods). (D) Snapshot of the TM domain structure at the end of the simulation is shown. The TM monomers are shown in same colors used in (B) and (C), and the phosphorus atoms of the lipid bilayer are shown as orange spheres.

**Figure S6:**
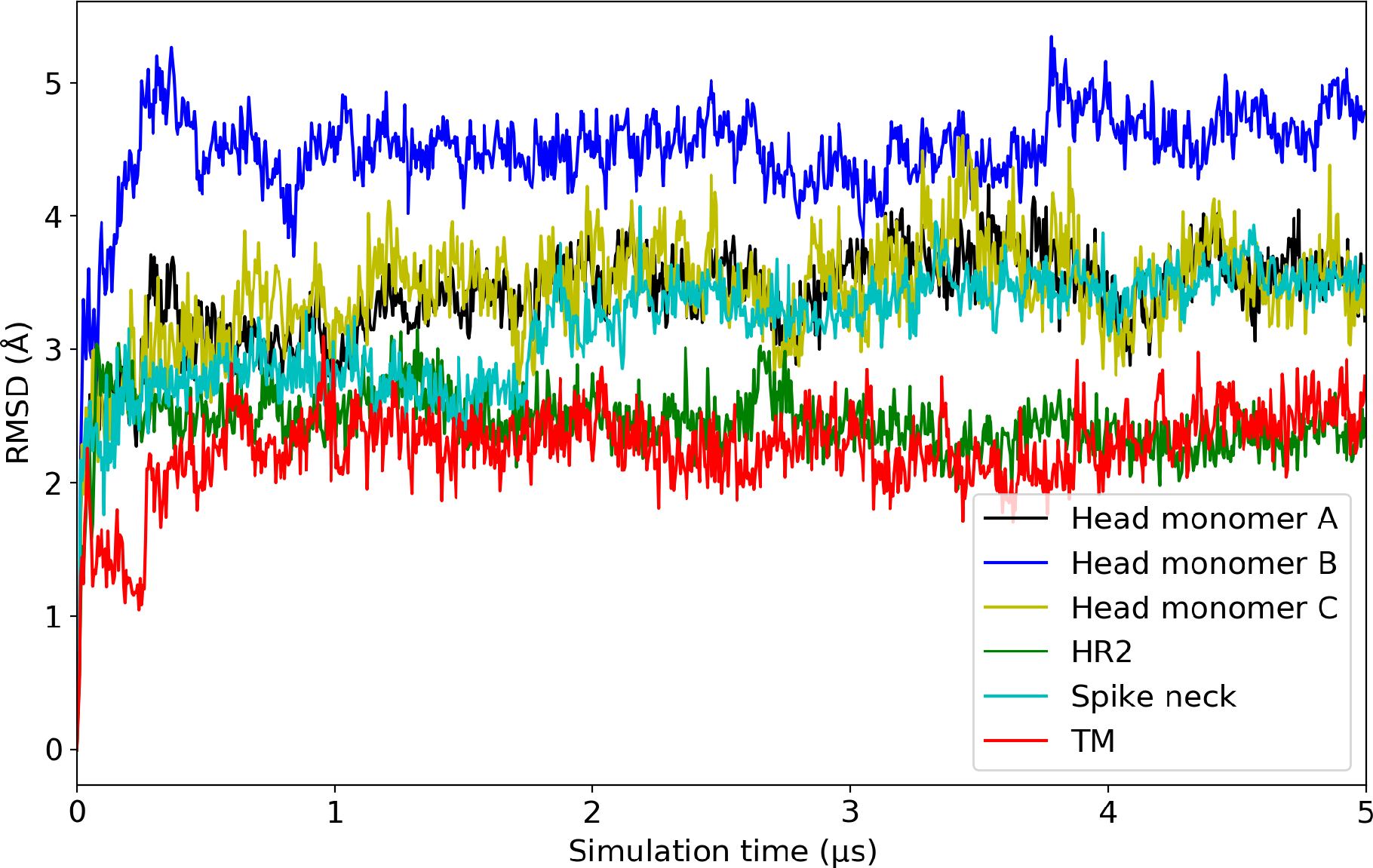
*C_α_* **RMSD of different spike domains:** The RMSD is calculated individually for different stalk domains: trimeric spike neck (residues 1139-1158), HR2 (residues 1173-1200), and the TM (residues 1213-1238), as well as the individual spike head monomers (monomer A, B and C (residues 1-1134). Monomer B of the spike head contains the RBD in the ”up” configuration and is relatively more mobile compared to the other monomers containing RBD in the ”down” configuration.

**Figure S7:**
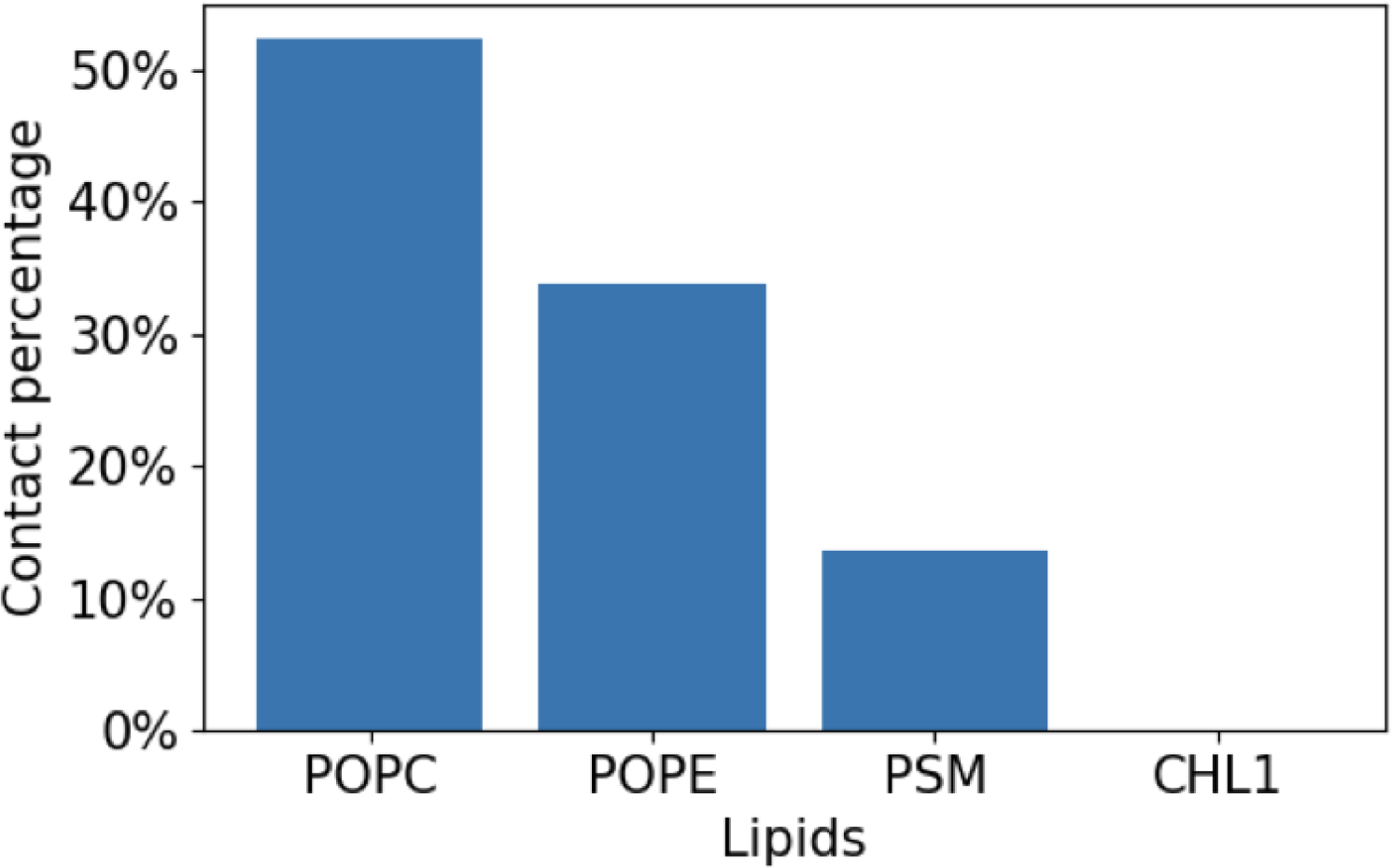
Lipid-glycan contacts with different lipid types. The contact frequency of different lipid types with the glycans at N1198 is shown. The reported percentages are normalized by the amount of POPC, POPE, PSM, and cholesterol in the leaflet proximal to the glycans.

**Figure S8:**
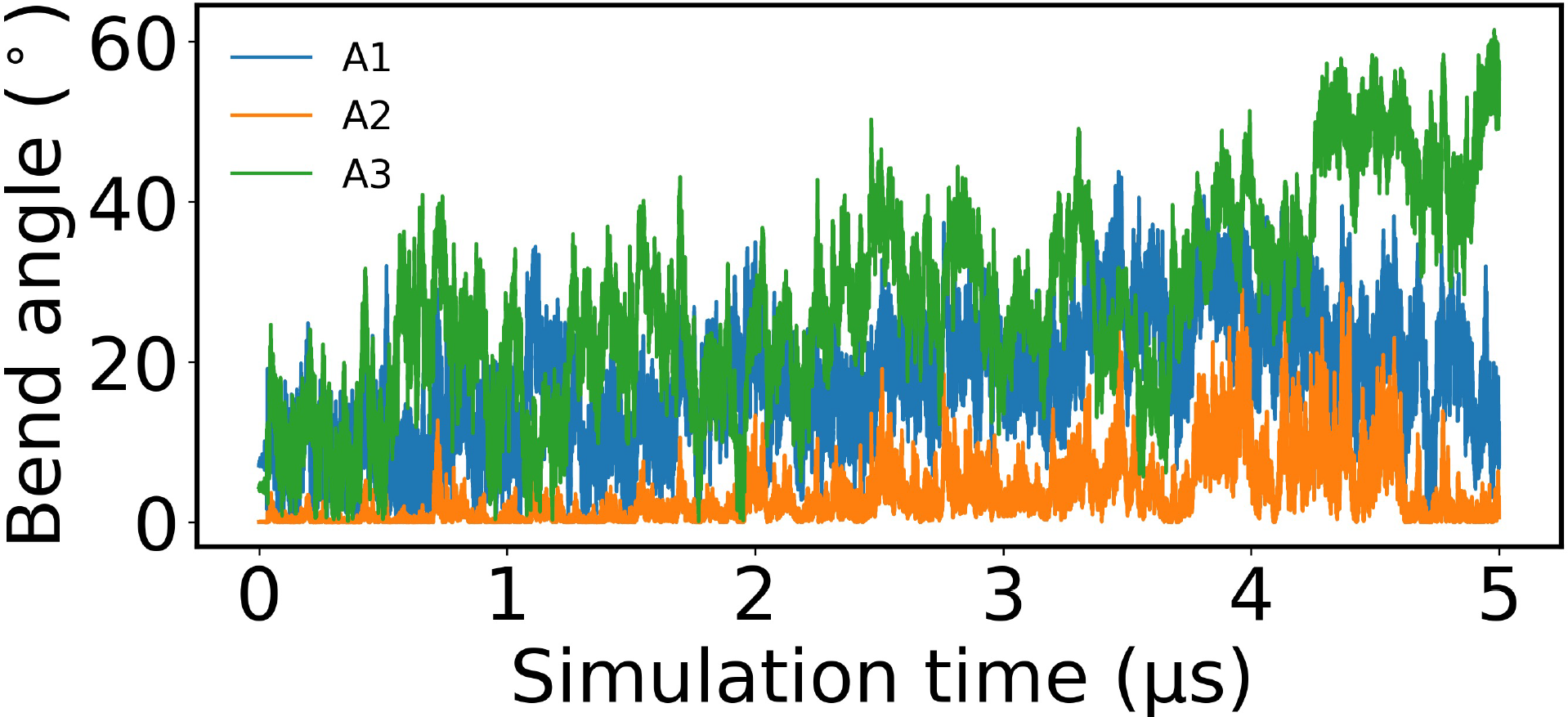
Correlation of the global motions in non-glycosylated spike. The time series of the A1, A2, and A3 angles depicted in Fig. 6A are shown for the simulation of the non-glycosylated spike.

**Figure S9:**
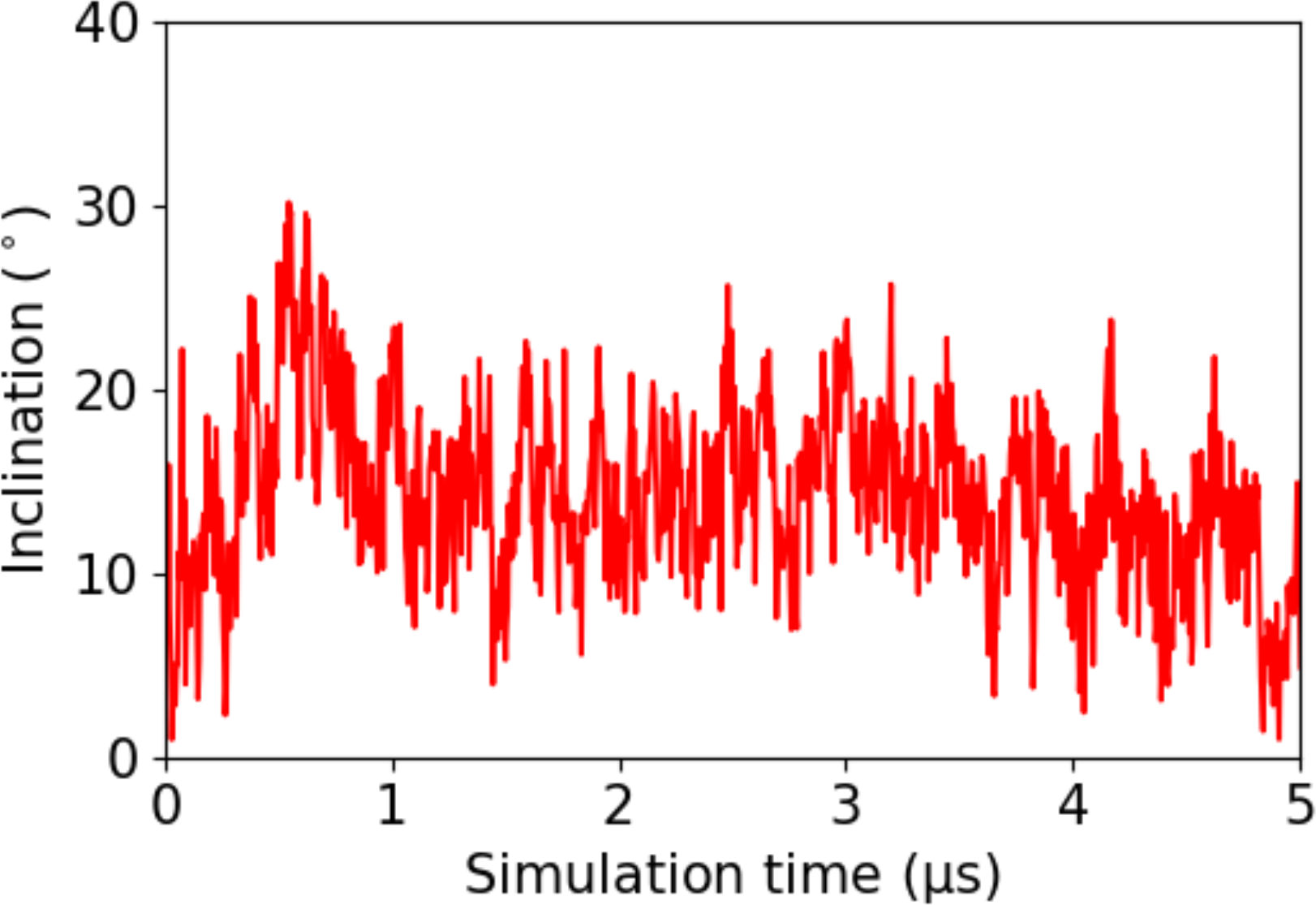
Tilt angle of the TM domain in the membrane. The tilt angle of the TM domain is calculated with respect to its initial orientation. The angle was measured between the third principal component of inertia calculated using all heavy atoms of the TM domain with respect to that for the starting structure.

**Table S1:** Lipid composition of the simulated membranes. The lipid composition (%) of the membrane patch representing the ERGIC lipid composition.

| **Leflet** | **POPC** | **POPE** | **Cholesterol** | **POPS** | **POPI** | **PSM** |
| --- | --- | --- | --- | --- | --- | --- |
| Outer | 59.0 | 25.5 | 15.0 | 0.0 | 0.0 | 0.5 |
| Inner | 45.0 | 18.5 | 15.0 | 6.0 | 15.0 | 0.5 |

**Table S2:** Glycosylation sites and types. The glycosylation sites in spike, as well as the respective composition of the sugar moiety. The sequence graph shown for the glycans follows the standard of Symbol Nomenclature For Glycans (SNFG).^87^

| Site | Composition and type | Sequence |
| --- | --- | --- |
| N61 | HexNAc(2)Hex(5)<br>M5 |  |
| N122 |  |  |
| N603 |  |  |
| N709 |  |  |
| N717 |  |  |
| N801 |  |  |
| N1074 |  |  |
| N234 | HexNAc(2)Hex(8)<br>M8 |  |
| N657 | HexNAc(3)Hex(6)<br>Hybrid |  |
| N149 | HexNAc(4)Hex(3)<br>Fuc(1)<br>Complex |  |
| N331 |  |  |
| N343 |  |  |
| N616 |  |  |
| N1134 |  |  |
| N1158 | HexNAc(4)Hex(4)<br>Complex |  |
| N1098 | HexNAc(4)Hex(4)<br>Fuc(1)NeuAc(1)<br>Complex |  |
| N165 | HexNAc(4)Hex(5)<br>Fuc(3)NeuAc(1)<br>Complex |  |
| N282 | HexNAc(5)Hex(3)<br>Fuc(1)<br>Complex |  |
| N17 | HexNAc(5)Hex(4)<br>Fuc(1)<br>Complex |  |
| N1173 | HexNAc(6)Hex(3)<br>Fuc(1)<br>Complex |  |
| N74 | HexNAc(6)Hex(4)<br>Fuc(1)NeuAc(1)<br>Complex |  |
| N1194 | HexNAc(6)Hex(5)<br>Fuc(1)NeuAc(1)<br>Complex |  |
| T323 | O-glycan |  |

**Table S3:**
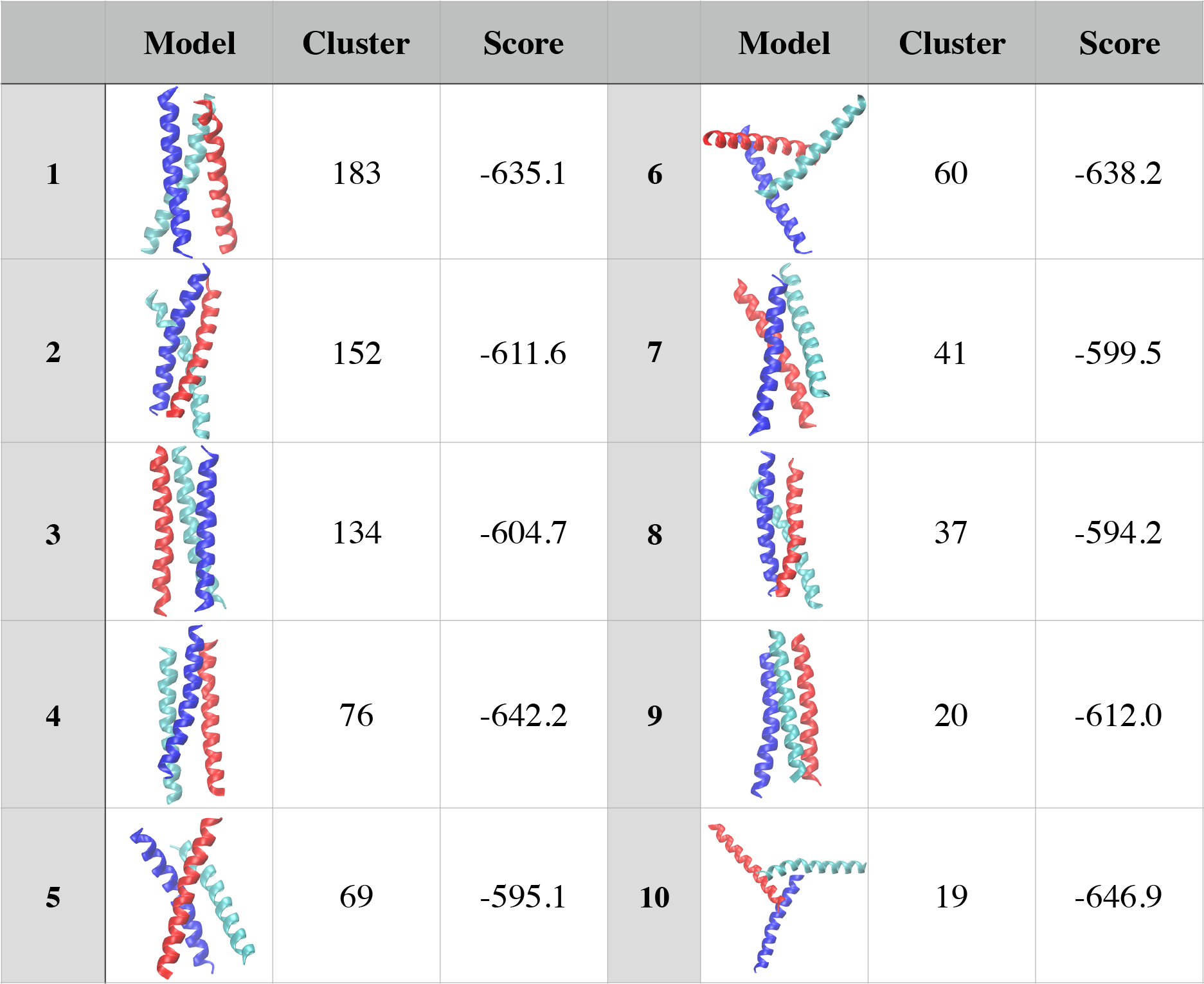
TM domain trimer models predicted by Cluspro. The predicted structures of the TM domain trimers as predicted by Cluspro^55^ are shown. The ’cluster’ column denotes the cluster size, which is the respective number of models present in each cluster after clustering is carried out on the selected TM domain trimer models. The ’score’ shows the predicted binding energy calculated using the energy function of Cluspro.

|  | Model | Cluster | Score |  | Model | Cluster | Score |
| --- | --- | --- | --- | --- | --- | --- | --- |
| **1** |  | 183 | -635.1 | **6** |  | 60 | -638.2 |
| **2** |  | 152 | -611.6 | **7** |  | 41 | -599.5 |
| **3** |  | 134 | -604.7 | **8** |  | 37 | -594.2 |
| **4** |  | 76 | -642.2 | **9** |  | 20 | -612.0 |
| **5** |  | 69 | -595.1 | **10** |  | 19 | -646.9 |

